# Impacts of *Desulfobacterales* and *Chromatiales* on sulfate reduction in the subtropical mangrove ecosystem as revealed by SMDB analysis

**DOI:** 10.1101/2020.08.16.252635

**Authors:** Shuming Mo, Jinhui Li, Bin Li, Ran Yu, Shiqing Nie, Zufan Zhang, Jianping Liao, Qiong Jiang, Bing Yan, Chengjian Jiang

## Abstract

Sulfate reduction is an important process in the sulfur cycle. However, the relationship between this process and the genotype of microorganisms involved in subtropical mangrove ecosystems is poorly understood. A dedicated and efficient gene integration database of sulfur metabolism has not been established yet. In this study, a sulfur metabolism gene integrative database (SMDB) had been constructed successfully. The database achieved high coverage, fast retrieval, and low false positives. Then the sulfate reduction by microorganisms in subtropical mangroves ecosystem had been evaluated quickly and accurately with SMDB database. Due to the environmental factors, dissimilatory sulfite reductase was significantly high in mangrove samples. Sulfide, Fe, and available sulfur were found to be the key environmental factors that influence the dissimilatory sulfate reduction, in which sulfur compounds could be utilized as potential energy sources for microorganism. Taxonomic assignment of dissimilatory sulfate-reduction genes revealed that *Desulfobacterales* and *Chromatiales* are completely responsible for this process. Sulfite reductase can help the community cope with the toxic sulfite produced by these Bacteria order. Collectively, these findings demonstrated that *Desulfobacterales* and *Chromatiales* play essential roles in dissimilatory sulfate reduction.

## 1 Introduction

Dissimilatory sulfate reduction, which results in the conversion of sulfate (SO_4_^2-^) to of HS^-^or H_2_S, is an important reaction in the sulfur cycle (Wenk et al., 2018). The study of dissimilatory sulfate reduction can reveal the occurrence of all dissimilatory sulfate-reduction genes in a community. However, sulfate reduction lacks a complete pathway in single strains, a condition that might be a common occurrence (Bukhtiyarova et al., 2019). The high occurrence of partial sulfate reduction implies that environmental conditions can affect gene occurrence. Gene families, including adenylyl sulfate reductase (*sat*), adenylyl sulfate reductase (*aprAB*), and dissimilatory sulfite reductase (*dsrABC*), are involved in the canonical dissimilatory sulfate-reduction pathway (Jochum et al., 2018, Anantharaman et al., 2018). Recently, the gene family of dissimilatory sulfite reductase beta subunit (*dsrB)* has been applied to study the diversity of sulfate-reducing bacteria (SRB) (Vavourakis et al., 2019). Dissimilatory sulfate reduction is primarily driven by SRB, and a complete absence of oxygen (O_2_) or low oxygen condition is vital for SRB to gain energy (Wu et al., 2017).

To the best of our knowledge, a dedicated sulfur database has not been constructed yet. Different databases, such as Clusters of Orthologous Groups (COG) (Galperin et al., 2015), Kyoto Encyclopedia of Genes and Genomes (KEGG) (Kanehisa et al., 2016), Evolutionary Genealogy of Genes: Non-supervised Orthologous Groups (eggNOG) (Huerta-Cepas et al., 2016), SEED subsystems (Overbeek et al., 2005), and M5nr (Wilke et al., 2012), are widely used for functional assignment of predicted genes. However, the application of these databases in analyzing sulfur cycle via shotgun metagenome sequencing data has several obstacles. First, few of these databases contain a complete database of gene families involved in sulfur cycle. Second, the use of these databases obtains a wrong relative abundance of gene families because of similar functional domains of the genes. For example, certain gene families, such as polysulfide reductase chain A (*psrA)* and thiosulfate reductase (*phsA)*, share high sequence similarity (Stoffels et al., 2012). This condition often leads to the mismatch interpretation of potential ecological processes. Finally, searching for sulfur cycle genes in large databases is time consuming. Therefore, a specific database of genes related to the sulfur cycle pathway should be established to address these problems. This novel database can also be applied to shotgun metagenomics research.

Mangrove sediments are usually characterized as anoxic with high levels of sulfur and salt and rich in nutrients (Ferreira et al., 2010). Sulfate reduction is one of the most active processes occurring in mangrove ecosystem (Lin et al., 2019). Dissimilatory sulfate reduction drives the formation of enormous quantities of reducing sulfide. The study of sulfate reduction in mangroves is interesting and important, but this endeavor has several limitations. First, although the diversity of sulfate-reducing bacteria has been studied in mangrove ecosystems, an understanding of sulfate reduction in these ecosystems remains insufficient (Wu et al., 2019). Second, researchers have studied culturable sulfate reduction of single strains via genomic analysis in other ecological environments (Spring et al., 2019), but it was not well study in mangrove ecosystems. Finally, the relationship between sulfate reduction and the genotype of microorganisms involved in this process in mangroves are also poorly understood. Furthermore, the environmental conditions that select for dissimilatory sulfate-reduction gene families for a more frequent reliance on sulfate reduction remain unclear.

Therefore, in this study, we created a manually managed database that primarily gathers most of the publicly available sulfur cycle gene families to address the limitations of available public databases in sulfur cycle analysis. An integrated database, namely, sulfur metabolism gene integrative database (SMDB), covering sulfur cycle gene families was established. SMDB was applied to samples from the subtropical mangrove ecosystem of Beibu Gulf in China to generate functional profiles. And then the sulfate reduction in the mangrove ecosystem had been evaluated. This study presents unique microbial features as a consequence of adapting to distinct environmental conditions to dissimilatory sulfate-reduction process.

## 2 Materials and methods

### 2.1 Sulfur metabolism gene integrative database development

A manually created integrated database was constructed to profile sulfur cycle genes from shotgun metagenomes, as the described with Tu et al. (2019), with slight modifications. The database was expected to bear the following characteristics: all sulfur gene families should be accurate and false positives should be minimized in database searching. The framework is shown in Supplementary Fig. S1.

#### 2.1.1 Sulfur metabolism gene integrative database sources

The universal protein (UniProt) database (http://www.uniprot.org/) was utilized to retrieve core database sequences of sulfur cycle gene families by using keywords. Other databases were used for retrieving nontarget homologous sequences. The other databases included COG (ftp://ftp.ncbi.nih.gov/pub/COG/COG2014/data), eggNOG (http://eggnogdb.embl.de/download/eggnog_4.5/), KEGG (http://www.genome.jp/kegg/), and M5nr (ftp://ftp.metagenomics.anl.gov/data/M5nr/current/M5nr.gz). In addition, the homologous sequences from RefSeq non-redundant proteins (NR) database (ftp://ftp.ncbi.nlm.nih.gov/blast/db/) were also added in the core constructed database. These databases were selected because they are widely used in metagenome research.

#### 2.1.2 Core database for constructing sulfur cycle genes

The KEGG and MetaCyc databases (Caspi et al., 2016) were referenced to manually retrieve gene families involved in sulfur cycle. Only those genes that had been experimentally confirmed to be involved in sulfur metabolism were collected through an extensive search of the literature. This step was followed by searching the Uniprot database with their keywords to identify the corresponding annotated sulfur metabolism gene families. For gene families with vague definitions (e.g., *cysQ* and *MET17*), the full protein name was added in the keywords to exclude sequences containing vague annotations. These datasets were retained as the core database for sulfur cycle genes.

#### 2.1.3 Full sulfur metabolism gene integrative database construction

To reduce false positives in database searching and increase the comprehensiveness of the core database, COG, eggNOG, KEGG, M5nr, and NR databases were used to retrieve nontarget homologous sequences to construct a full database. The full sulfur metabolism gene integrative database included the core database of sulfur cycle gene families and their nontarget homologous gene families. The sequence files of this integrative database were clustered using the CD-HIT program (version 4.6) (Fu et al., 2012) at 95% and 100% identity cutoffs. Representative sequences from this integrative database were then selected to construct SMDB. SMDB database had been deposited in https://github.com/taylor19891213/sulfur-metabolism-gene-database on January 8th, 2020. The SMDB website (http://smdb.gxu.edu.cn/) had been online since June 22th, 2020.

### 2.2 Case study

#### 2.2.1 Sampling sites and sediment collection

The ShanKou mangrove sediment in Beihai City, China (21°29′25.74″N, 109°45′49.43″E) was selected as the sampling location (Supplementary Fig. S4). All the sediment samples were collected on March 27, 2019. At each sample point (0-5 cm), triplicates sediment within an area of 10 m × 10 m were randomly collected using polyvinyl chloride tubes with a diameter of 3 cm and a height of 15 cm. Samples were placed in a sterile bag inside a box filled with ice immediately after sufficient mixing. All samples were stored at -80 °C in the same day. Rhizosphere samples (RS) from near-root were designated as RS1, RS2, and RS3. Nonrhizosphere samples (NRS) from nonrhizosphere were labelled as NRS1, NRS2, and NRS3. Nonmangrove samples (MS) were assigned as NMS1, NMS2, and NMS3. Mangrove samples (MS) included groups of RS and NRS. Sediment samples were air-dried and sieved through 0.25 mm steel mesh to remove stones and coarse materials, used for the chemical analysis.

Several sediment properties, such as total organic carbon (TOC), total organic nitrogen (TN) content, nutrients (NH_4_^+^, NO_3_^-^), and available sulfur (AS), were determined. Redox potential (ORP) was measured in situ by using a portable ORP meter (BPH-220, Bell, China). Soil suspension was obtained by centrifugation to determine pH by using a pH meter (PHS-2C; Sanxin, China) and salinity by using a salinity meter (PAL-06S; Atago, Japan). Available sulfur was determined by Barium sulfate turbidimetry. Available sulfur includes soluble sulfur, absorbent sulfur and partly of organic sulfur. Total nitrogen (TN) was determined by acid fumigation method and followed the method described by Harris et al. (2001). TP was determined by alkali digestion method and followed the method described by Yang et al. (2013). Fe concentration was determined by Atomic Absorption Spectrophotometer (GBC932, Australia) according to the instruction. Sulfide was determined by methylene blue spectrophotometric. TOC were determined with an Automatic Carbon and Nitrogen Analyzer (TOC-TN 1200 The TOC were rmo Euroglas) according to the instruction. A detailed description of the analytical methods used for NH_4_^+^ and NO_3_^-^, was previously published (Yang et al., 2017).

### 2.2.2 DNA extraction and high-throughput sequencing

DNA was extracted from the sediments by using the FastDNA SPIN kit for soil (MP Biomedicals, USA) according to the manufacturer’s instructions. At least 6 μg DNA from each sample was submitted to the Shanghai Majorbio Bio-Pharm Technology Co., Ltd. (Shanghai, China) for sequencing on an Illumina platform. The data output from each DNA sample was over 10 Gb. The metagenomic sequencing data were deposited to the NCBI SRA database under the BioProject PRJCA002311 (https://bigd.big.ac.cn/gsub/).

### 2.2.3 Shotgun metagenomic sequence processing and analysis

Initial quality assurance/quality control, including trimming sequencing adapters and bar codes from sequence reads, was performed. Adapter sequences were removed using SeqPrep v1.33 (https://github.com/jstjohn/SeqPrep). Moreover, sequences less than 100 bp, sequences with quality of less than 20, and reads containing an N base were removed using Sickle v1.2 (Joshi NA, 2011). Finally, clean reads were created. Sequences were assembled using megahit (v1.1.3) with the default parameters (Li et al., 2016). Open reading frames (ORFs) were predicted translated by using Prodigal (v3.02) (Hyatt et al., 2010) and unigenes were identified with CD-HIT (v4.5.6) (Fu et al., 2012). For unigenes, relative abundance is calculated based on the length of genes with same genes assignment in the sum of reads.

For functional annotation, unigenes were aligned against the online SMDB database website (http://smdb.gxu.edu.cn/). The blast software of the SMDB database is DIAMOND v0.9.14 (Buchfink et al., 2015) with parameters set as an e-value cutoff of 1 × 10^−5^ by using BLASTP. The online website of the SMDB database output was converted to m8 blast format. Gene families of dissimilatory sulfate reduction were screened out. The best hits were extracted for sulfur cycle genes profiling. A correlation heat map was used to visualize the composition of dissimilatory sulfate reduction across all nine samples. Spearman correlation coefficients of abiotic factors and dissimilatory sulfate-reduction genes were calculated by SPSS (George and Mallery, 2013).

For dissimilatory sulfate reduction genes, their taxonomy was assigned based on BLASP results in the NCBI NR database with coverage > 50% and e value < 1 × 10^−10^. The methods for gene taxonomy as Hou et al. described (Hou et al., 2020). Taxonomic assignment was summarized at the order level. For each functional gene, taxonomic relative abundance is calculated based on the sum sequencing depth of genes with same taxonomic assignment in the total depth of this gene.

### 2.2.4 Annotation of taxonomy

For the taxonomic annotation, the unigenes were aligned against NR database using a local blastp program (E-value set as 10^−10^). A blastp result was imported into MEGAN Community, and taxonomic classification was performed using LCA algorithm (Huson et al., 2016).

## 3 Results

### 3.1 Characteristics of sulfur metabolism gene integrative database

A total of 175 gene (sub)families covering 11 sulfur metabolism pathways, including assimilatory sulfate reduction, thiosulfate disproportionation, sulfide oxidation, dissimilatory sulfate reduction, sulfite oxidation, sulfur oxidation, sulfur reduction, tetrathionate oxidation, tetrathionate reduction, thiosulfate oxidation, and organic degradation/synthesis, were recruited in SMDB (Supplementary Table S1). SMDB obtained 115,321 and 395,737 representative sequences at 95% and 100% identity cutoffs, respectively.

To demonstrate the necessity of building a manually created sulfur metabolism gene database, we first compared the coverage and accuracy of sulfur metabolism gene (sub)families in SMDB with those of existing public databases. Among the 175 gene (sub)families we recruited in SMDB, only 118, 157, 144, 169, and 172 could be found in these gene (sub)families in COG, eggNOG, KEGG, M5nr, and NR databases, respectively (Fig. 1A, Supplementary Table S1). The accuracy of SMDB containing sulfur gene (sub)families exceeded that of public databases (Supplementary Fig. S2). We characterized sulfur metabolism gene (sub)families by searching nine shotgun metagenomic datasets obtained by sampling the mangrove environment against SMDB, COG, eggNOG, KEGG, M5nr, and NR databases. Subsequently, the number of sulfur metabolism gene families detected by searching against SMDB far exceeded that of the five other databases (Supplementary Fig. S3).

**Figure 1.**
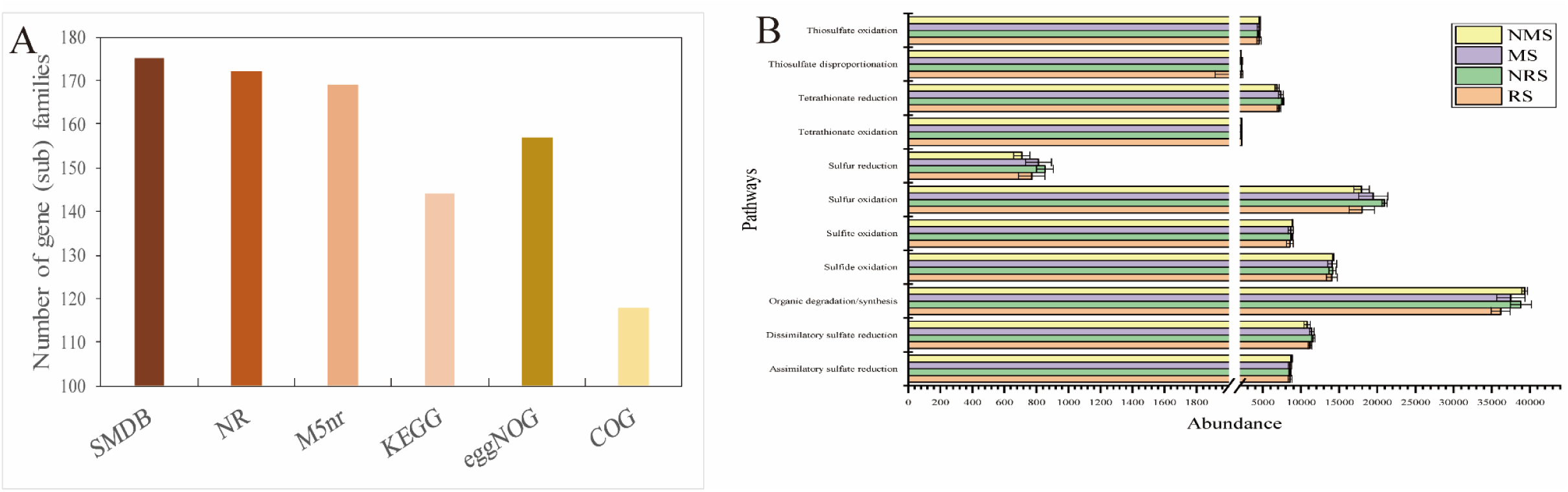
A) Sulfur cycle gene (sub)families were detected using SMDB, NR, M5nr, KEGG, eggNOG, COG databases. B) Pathway abundances in the samples. The Y-axis is gene (sub)families number in different databases.

### 3.2 Abundance and diversity of sulfur (sub) gene families

A total of 151 different sulfur gene (sub)families were annotated. The sulfur gene (sub)families in each sample ranged from 142 (sample RS3) to 146 (sample RS3) (Supplementary Table S2). The abundance of pathways showed that the organic sulfur degradation/synthesis pathway in these samples was the highest, followed by sulfur oxidation; the pathway with the lowest abundance was sulfur reduction except for others (Fig. 1B). As shown in Supplementary Table S2, the most abundant genes were sulfonate transport system ATP-binding protein (*ssuB)*, heterodisulfide reductase (*hdrA/D*), arylsulfatase (*atsA*), and dimethylsulfoniopropionate demethylase (*dmdB/A*).

### 3.3 Microbial diversity based on metagenome

To study the distribution of dissimilatory sulfate reduction in microbial communities, we annotation the taxonomy. The taxonomic assignments indicated that members of *Desulfobacterales* was dominant, with compositions ranging from 16% to 22%, across each sample (Fig. 2A). Other order, such as *Spirochaetales, Cellvibrionales*, and *Gemmatimonadales* comprised apprximately 4% (Fig. 2A). The abundance of *Desulfatibacillum, Desulfobacterium*, and *Desulfosarcina* in MS exceeded that in NMS, while *Desulfobacter, Desulfobulbus*, and *Desulfurivibrio* in RS exceeded that in NRS (Fig. 2B).

**Figure 2.**
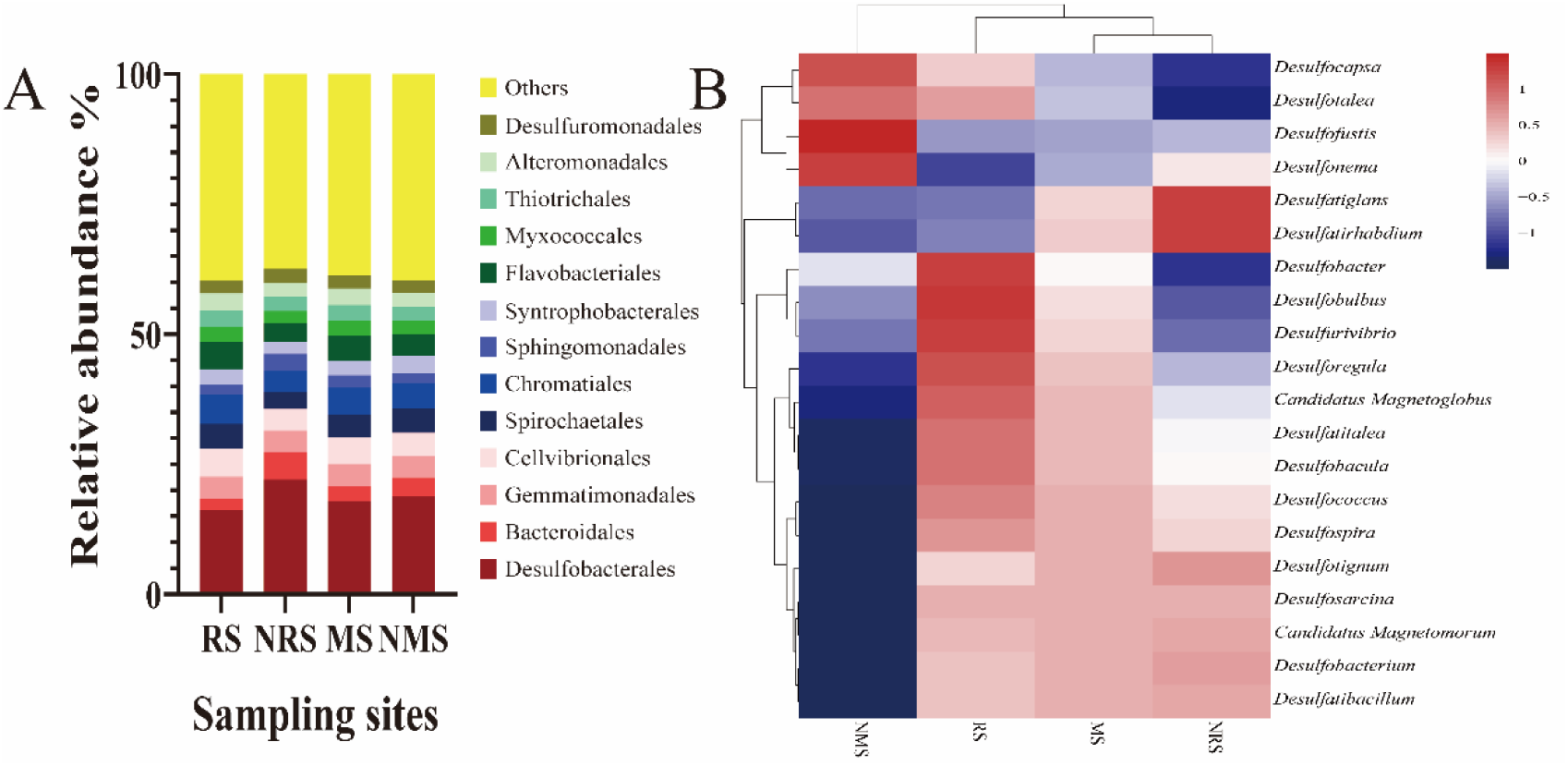
Comparison of taxonomic assignments from the metagenomes. A) The 13 dominant order are shown with their relative abundances; the remaining order are indicated as “Others”. B) The heat map of *Desulfobacterales* order.

### 3.4 Genes for dissimilatory sulfate reduction

The prevalence of dissimilatory sulfate-reduction genes at the sites was initially analyzed by considering the complete dissimilatory sulfate-reduction pathway. Accordingly, genes encoding adenylyl sulfate transferase (*sat*), adenylyl sulfate reductase (*aprA/B*), and dissimilatory sulfite reductase (*dsrA/B*) were obtained (Fig.3A). In the MS and NMS groups, *dsrA,aprA* and sulfite reduction-associated complex DsrMKJOP multiheme protein (*dsrM/P*) were significantly high in mangrove samples (p < 0.05, Fig. 3B). While, genes encoding sulfate adenylyltransferase subunit 2 (*cysD*) was significantly low in mangrove samples. *DsrA* catalyzes the reduction of sulfite to sulfide. This is the terminal oxidation reaction in sulfate respiration (Pott et al., 1998). *AprA* which catalyzes reversibly the reduction of adenosine 5’-phosphosulfate (APS) to sulfite and AMP during dissimilatory sulfate reduction (Chiang et al., 2009). In the RS and NRS groups, genes encoding anaerobic sulfite reductase (*asrB*) and *DsrC*-disulfide reductase (*dsrK*) were significantly low in rhizosphere samples (p < 0.05, Fig. 3C). Genes *sat* and *aprA* were more frequent than *dsrA/B* (Fig. 3A). The gene of *sat*, accounting for 9.6%–13.3% of the total dissimilatory sulfate-reduction genes in all samples (Fig. 3A).

**Figure 3.**
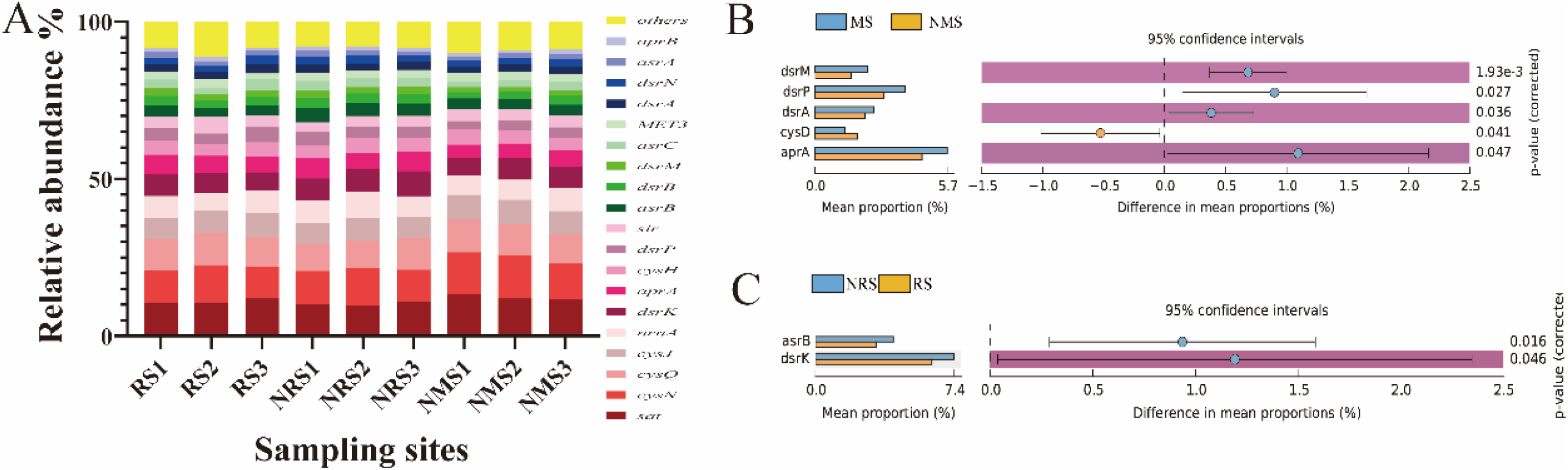
A) The bar chart indicates the relative abundances of the 20 abundant dissimilatory sulfate reduction gene families in each sample. Significant differences in sulfur (sub) gene families between mangrove and nonmangrove sediments (B) and rhizosphere and nonrhizosphere sediments (C).

### 3.5 Taxonomic assignment of dissimilatory sulfate-reduction genes

*Desulfobacterales* were the predominant in microbial community. Therefore, taxonomic assignments were done at the order level to obtain good process of dissimilatory sulfate-reduction genes families (Table 1). The numbers of dissimilatory sulfate-reduction genes form *Desulfobacterales* and *Chromatiales* exceeded those of other taxonomic assignments, whereas the remaining genes were assigned to *Rhizobiales, Desulfovibrionales, Desulfuromonadales* and *Cellvibrionales* (Supplementary Fig. S5). This finding was consistent with the microbial diversity analysis showed that these samples were as prevalent as Desulfobacterales (phylum *Proteobacteria*) (Fig. 2A). The high abundance of *aprA* was obtained from *Desulfobacterales* and *Chromatiales* (Table 1). In addition, *asr* and *cys* gene families were obtained from *Desulfobacterales* and *Chromatiales* (Table 1). Taxonomic classification of *dsrB* was assigned to *Chromatiales* (11.14%) and *Desulfobacterales* (67.26%) in MS, while it was assigned to *Chromatiales* (22.66%) and *Desulfobacterales* (48.50%) in NMS (Supplementary Fig. S7). However, taxonomic classification of *dsrB* was assigned to *Chromatiales* (9.10%) and *Desulfobacterales* (65.72%) in RS, while it was assigned to *Chromatiales* (12.44%) and *Desulfobacterales* (68.23%) in NRS (Supplementary Fig. S8).

**Table 1.** Taxonomic assignments of dissimilatory sulfate-reduction genes.

| taxonomy | RS | NRS | NMS |
| --- | --- | --- | --- |
| *Chromatiales* | *cysJ; dsrK; aprA; cysN; dsrA;*<br>*sir; sat; aprM; cysQ; cysNC;*<br>*cysD; cysH; dsrO; asrB; dsrN;*<br>*dsrP; asrA; nrnA; MET3; dsrB;*<br>*asrC; HINT4; cysI* | *cysN; aprA; dsrB; dsrO; sir;*<br>*sat; cysQ; dsrK; cysNC; cysD;*<br>*MET3; dsrM; cysH; cysJ; asrB;*<br>*dsrP; dsrN; cysI; dsrA; aprB;*<br>*nrnA; dsrC; HINT4* | *cysJ; cysN; asrB; cysD; cysNC;*<br>*dsrK; asrA; sat; cysI; cysH; cysQ;*<br>*aprA; dsrC; sir; aprM; nrnA; dsrP;*<br>*MET3; dsrM; aprB; dsrB; dsrA* |
| *Desulfobacterales* | *cysJ; cysN; aprA; asrB; cysQ;*<br>*asrC; dsrK; nrnA; sat; dsrM;*<br>*asrA; dsrN; cysH; dsrB; sir;*<br>*dsrP; cysD; SAL; MET3; cysNC;*<br>*dsrA; aprB; cysI; dsrO; MET22;*<br>*dsrC; dsrJ* | *dsrK; cysJ; sat; aprA; cysN;*<br>*cysQ; nrnA; dsrJ; dsrM; dsrC;*<br>*asrB; cysNC; dsrN; cysH; dsrB;*<br>*asrC; dsrA; sir; dsrP; aprB; SAL;*<br>*asrA; MET3; MET22; dsrO; cysI;*<br>*cysD* | *dsrK; cysJ; sat; asrB; cysN; asrA;*<br>*cysQ; aprA; dsrB; MET22; nrnA;*<br>*sir; dsrP; aprB; dsrJ; dsrN; dsrM;*<br>*cysD; cysNC; dsrA; asrC; cysH;*<br>*dsrO; cysI; MET3; dsrC* |
| *Rhizobiales* | *sat; dsrO; cysI; cysJ; cysH;*<br>*asrA; cysQ; cysN; sir; nrnA;*<br>*asrB* | *sat; aprA; cysJ; cysH; cysQ;*<br>*asrB; HINT4; cysN* | *aprM; sat; dsrA; aprA; cysN; cysI;*<br>*cysQ; asrB; cysJ; dsrO; sir; dsrK;*<br>*nrnA* |
| *Desulfovibrionales* | *sat; asrC; dsrK; sir; cysN; cysJ* | *aprA; sat; dsrJ; cysN; asrC; dsrK; cysJ* | *sat; dsrK; cysN* |
| *Desulfuromonadales* | *cysI; cysN; asrB; nrnA; cysQ; sir;*<br>*asrC* | *cysN; sir; asrC* | *cysI; cysN; asrB; sat; dsrA; sir; cysQ;*<br>*dsrP; nrnA* |

### 3.6 Sediment properties

To determine the environmental factors that likely shape the structure and composition of dissimilatory sulfate-reduction genes in mangrove sediments, we generated a correlation heat map (Fig. 4). Certain sediment properties, such as pH, TOC, AS, ORP, NH_4_^+^, NO_3_^-^, TN, TP, Fe, salinity, and sulfide content, were determined (Supplementary Fig. S6). The concentrations of AS, Fe, TOC, and TN were significantly higher in the mangrove samples than those in the nonmangrove samples. Moreover, the concentrations of TOC, TN, and TP were significantly higher in the rhizosphere samples than those in the nonrhizosphere samples (Supplementary Tables S5 and S6, p < 0.05).

**Figure 4.**
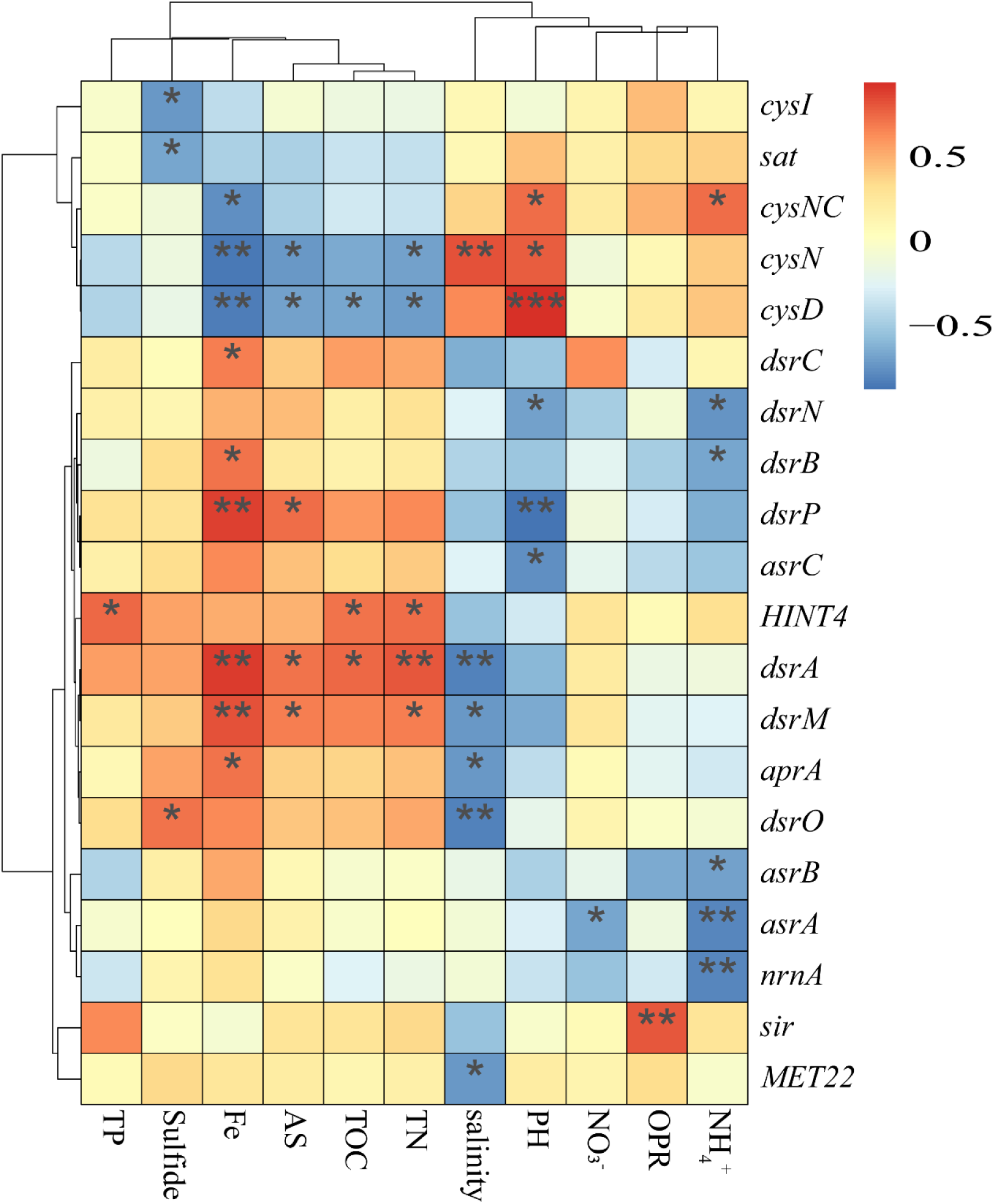
Correlation heat map according to z-scores of the 20 most abundant dissimilatory sulfate reduction gene (sub) families with significant correlation between sediment properties. * p < 0.05 ** p < 0.01

Result showed significant correlations between sediment properties and dissimilatory sulfate-reduction gene (sub) families (p < 0.05, Fig. 4, Supplementary Table S3). pH was negatively correlated with *cysD, dsrP*, and *asrB*. Salinity content was significantly correlated with *aprA*, and *dsrB/M*. Sulfide content was significantly correlated with *sat*, Sulfite reduction-associated complex DsrMKJOP (*dsrO*), and sulfite reductase (NADPH) hemoprotein beta-component (*cysI*). Fe content was significantly correlated with *dsrA/B/C/M*, and *aprA*. AS content was significantly correlated with *dsrA/M/P*. Results of Pearson correlation analysis of the pathways are summarized in Supplementary Table S4. Results showed no significant correlation between environmental factors and the dissimilatory sulfate-reduction pathway. AS was negatively correlated with the organic degradation/synthesis pathway. Fe was positively correlated with the sulfur reduction (Supplementary Table S4).

## 4 Discussion

Sulfur cycle is one of the most important biogeochemical cycles in Earth ecosystems. In this study, we developed a manually created database for the fast and accurate profiling of sulfur metabolism genes from shotgun metagenome sequencing data. The developed SMDB addresses the bottleneck encountered in the previous public databases. First, SMDB has a precise definition for sulfur metabolism gene families. Unlike other database generations (Huerta-Cepas et al., 2016), gene families in SMDB were manually retrieved by using keywords combined with sequence similarity. Typical examples include the *phsA* and *psrA* genes that encode particle thiosulfate reductase and polysulfide reductase, respectively, but these genes share high sequence similarity (Stoffels et al., 2012). Hence, distinguishing the activities of these two enzymes during genome annotation is difficult. Second, SMDB presents a comprehensive coverage of gene families in sulfur metabolism. We aimed to provide the latest knowledge and research progress in sulfur metabolism studies, such as aerobic dimethyl sulfide (DMS) degradation (Koch and Dahl, 2018). The gene families *TST/MPST/sseA* for K01011 display considerable sequence similarity in KEGG, but they are distinct in two sulfur transferase subfamilies. Finally, SMDB considers the problem of false positives. Herein, this issue was addressed by including homologous gene families from multiple public databases. Most importantly, the small size of SMDB reduces the computational cost required to output the functional profiling of files of sulfur metabolism gene families. The database achieved high coverage and accuracy (Supplementary Fig. S1). The discrepancy due to these databases have their own advantages as well as limitations owing to different designing concepts associated with each database. For example, the COG database mainly relies on complete microbial genomes (Galperin et al., 2015). The KEGG database holds advantages of linking genes to pathways in data interpretation (Kanehisa et al., 2016).

We applied SMDB in characterizing sulfur metabolism gene (sub)families in mangrove sediment environment to demonstrate its potential applications in microbial ecology studies. Most studies have used experiments for microbial. The activity of microbial metabolism can be determined by (radio-)tracer experiments. Polymerase chain reaction (PCR), a technique used to make numerous copies of a specific segment of DNA quickly and accurately. However, PCR usually produces bias, resulting in inaccurate experimental results because of lack of perfect working primers for many of the gene families involved (Peng et al., 2018). In the present study, shotgun metagenome sequencing technology was used to investigate the complete sulfur cycle in mangroves. The metagenomics survey described herein provided strong evidence that in neutral pH, anoxic, high-sulfur soil, the environment in the Beibu Gulf of China biases the dissimilatory sulfate-reduction genotype. AS and Fe concentrations were consistently higher in the mangrove zones than in the nonmangrove zones (p < 0.05, Supplementary Table S5), suggesting that the concentrations of AS and Fe determine the abundance of dissimilatory sulfate-reduction genes. Sulfide, Fe, and available sulfur were found to be the key environmental factors that influence the dissimilatory sulfate reduction genes (p < 0.05, Fig. 4). However, our current understanding of dissimilatory sulfate reduction suggests that Sulfide, Fe, and available sulfur contents are insufficient to explain gene trends, and available O_2_ is also an important factor. Mangrove sediments are usually characterized as anoxic (Ferreira et al., 2010). Mangrove conditions favoring dissimilatory sulfate reduction were probably of sufficient duration to contribute to the patterns observed herein, but a more detailed study of sediment properties should be conducted for confirmation.

This pattern supported the hypothesis that environmental conditions act on genes in the pathway and biases the dissimilatory sulfate-reduction genotype. In all samples, *aprA* occurred at higher levels than that from dissimilatory sulfate-reduction genes (*dsrA/B*) and anaerobic sulfite reductase (*asrA/B*) (Fig. 3A). This consistent with the previous studies, the occurrence of inorganic sulfur cycle genes in sequence genomes found that *aprA* occurs at higher frequencies than the other gene-encoding proteins required for dissimilatory sulfate-reduction genes (Vavourakis et al., 2019). This condition is probably due to the simultaneous sulfate reduction and sulfide oxidation processes (Watanabe et al., 2013). *DsrA* and *aprA* were significantly high in mangrove samples (p < 0.05, Fig. 3B). Owing to the unique ecological environments of subtropical mangrove ecosystems, the class *Proteobacteria* of *Betaproteobacteria* is the dominant SRB group in mangrove (Varon-Lopez et al., 2014). However, our results show that *Deltaproteobacteria* (order *Desulfobacterales, Chromatiales)* was the dominant SRB group in mangrove (Supplementary Fig. S5). In addition, *Desulfobacterales, Chromatiales* had relatively complete gene families of dissimilatory sulfate reduction in this study (Table 1). The *aprA* were basically obtained from *Desulfobacterales* and *Chromatiales* (Table 1, Supplementary Fig. S7), suggesting that the function of *aprA* in mangrove ecosystems may be mainly performed by these *Bacteria orde*r. Microbial sulfate reduction is important for anaerobic degradation of organic matter, and is often mediated by *Deltaproteobacteria*, specifically, *Desulfobacterales* (Meyer et al., 2007). Taxonomic classification of *dsrB* found that *Desulfobacterales* was high in MS than NMS. These findings showed that unique ecological environments selected these microbial communities for dissimilatory sulfate reduction.

Notably, sulfite is cytotoxic above a certain threshold if not rapidly metabolized and can wreak havoc at the cellular and whole-plant levels. By providing the gene families of *asrC, asrA, asrB, cysI, cysJ*, and *cysH* that reduces sulfite levels to protect the members of the local community (Bender et al., 2019). These activities are required for the biosynthesis of L-cysteine from sulfate (Zeghouf and Chemistry, 2000). If sulfite reductase is absent, most organisms in the community would be inhibited and grow slowly. The sediment properties (e.g., OPR, Fe) results in the release of sulfides from sediments (Shi et al., 2016). Other studies have shown that sulfides in mangrove sediments are eventually converted to L-cysteine (Lin et al., 2019).

We depicted the process of sulfur metabolism in mangrove sediments. Sulfur metabolism in mangrove sediments includes biological and abiotic processes (Fig. 5). For abiotic processes, Fe(III) is buried and acts as an oxidant for sulfide in deeper sediment layers where it partly binds the produced sulfide as iron sulfide (FeS) and pyrite (FeS_2_) (Wasmund et al., 2017). In this study, the concentration of Fe was consistently higher in the mangrove zones than that in the nonmangrove zones (p < 0.05, Supplementary Table S5). H_2_S and FeS can be oxidized into various oxidation states of sulfur. For biological processes, the role of microorganisms includes both oxidation and reduction of sulfur compounds. Dissimilatory sulfate reduction involves the reduction of sulfate to sulfide where the electron acceptor is sulfate, APS, sulfite, elemental sulfur, etc. By contrast, the reduced sulfide is assimilated in proteins and amino acids in assimilatory sulfate reduction (Fuentes-Lara et al., 2019). Sulfide oxidation by O_2_, NO_3_^-^, and Fe^3+^as electron acceptors produces sulfur, while Mn^4+^ as electron acceptors produces thiosulfate or sulfate. In addition, sulfur undergoes coupled oxidation and reduction of sulfur compounds (thiosulfate, sulfite, and sulfur) to sulfate and sulfide. DMS was derived from the degradation of organic sulfur (e.g., dimethylsulfoniopropionate). The dimethylsulfide (DMS) pathway provides a new link between the cycles of organic and inorganic sulfur compounds (Koch and Dahl, 2018).

**Figure 5.**
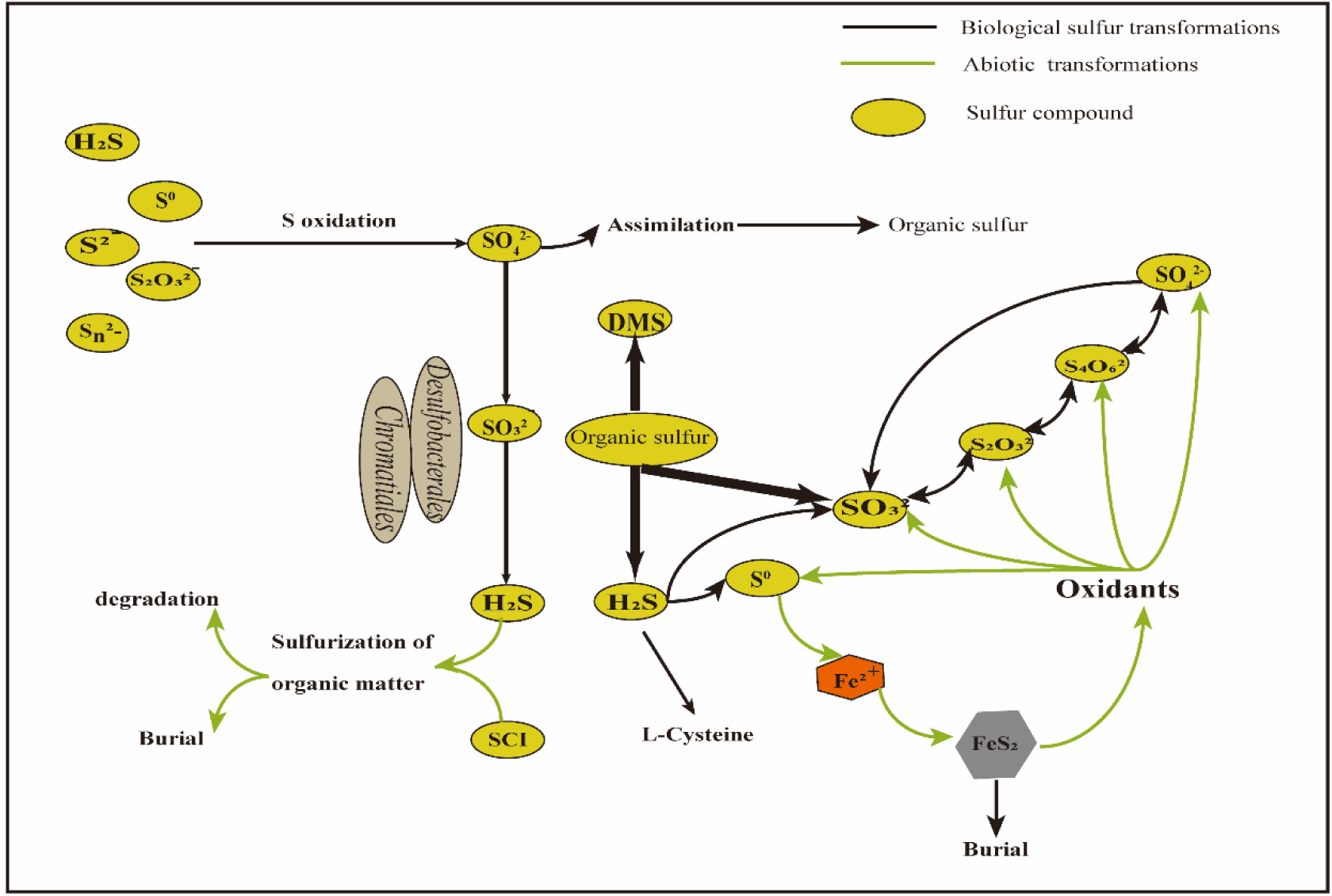
Conceptual depiction of sulfur metabolism in mangrove ecosystem, including reactions of inorganic and organic sulfur compounds and transformations of sulfur compounds of intermediate oxidation states (sulfur cycle intermediates, SCI). Inorganic sulfur compounds include oxidation and reduction of sulfur compounds. Black lines depict biological sulfur transformations by microorganisms. Green lines depict abiotic reaction-mediated sulfur transformations to pyrite (FeS_2_). Sulfur compounds are depicted within yellow eclipses.

The mechanism of dissimilatory sulfate-reduction metabolic pattern in mangrove ecosystem was constructed (Fig. 6). The figure shows the relative abundances of gene-encoding enzymes involved in dissimilatory sulfate reduction. The reduced sulfur compounds were likely oxidized to sulfite by reverse dissimilatory sulfite reductase (encoded by *dsrAB*) and thiosulfate sulfur transferase. EC 2.7.7.4 (sulfate adenylyltransferase) had the highest abundance among the enzymes. EC 1.8.99.2 (adenylyl-sulfate reductase) and EC 1.8.99.5 (dissimilatory sulfite reductase) had higher abundance in mangrove samples than in nonmangrove samples. The model of dissimilatory sulfite reductase also supports the hypothesis that mangrove samples prefer dissimilatory sulfate reduction. In rhizosphere samples, the abundance of EC 1.8.99.5 was low, whereas that of EC 1.8.1.2 (assimilatory sulfite reductase) was high. This finding indicated that rhizosphere microorganisms are conducive to the assimilation of sulfate to L-cysteine. Moreover, this result highlighted the importance of mangrove plants in sulfur cycle.

**Figure 6.**
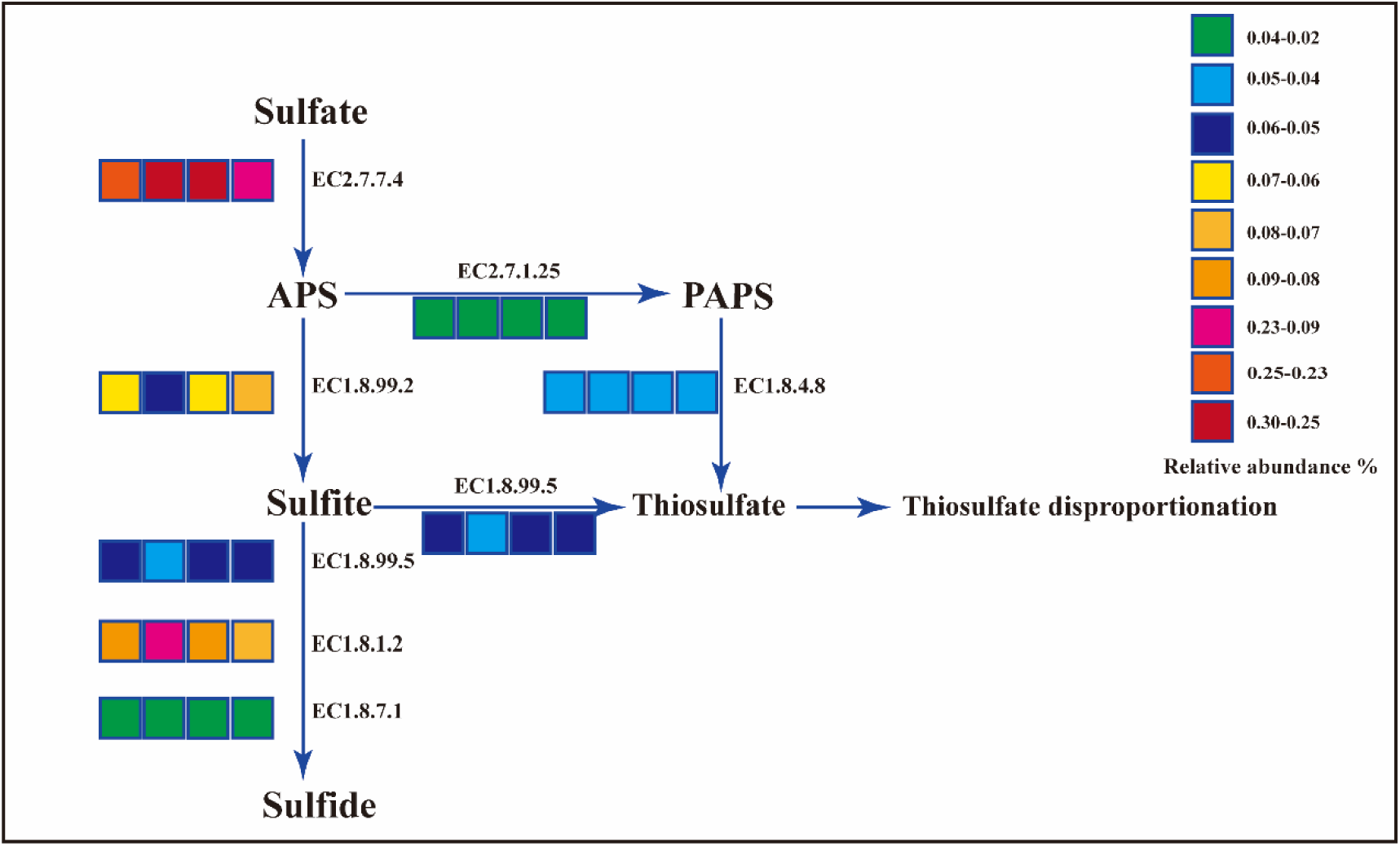
Model for the pathway of dissimilatory sulfate reduction. Relative abundances of the gene-encoding enzymes involved in dissimilatory sulfate reduction. The EC numbers of enzymes are boxed. The relative abundances of enzymes in the samples are shown in the nearby color bar, in which the four segments from left to right represent MS and NMS, RS, NRS, respectively.

## 5 Conclusions

In sum, a high-coverage database, namely, SMDB, was successfully constructed. This integrative database contains 175 gene (sub)families and covers 11 sulfur metabolism processes. SMDB was able to quickly and accurately analyze sulfur metabolism. The subtropical mangrove ecosystem biases dissimilatory sulfate-reduction genotype. Sulfide, Fe, and available sulfur were found to be the key environmental factors that influence the dissimilatory sulfate reduction. *Desulfobacterales* and *Chromatiales* were found to be responsible for dissimilatory sulfate reduction. This study provides a theoretical basis for the mechanism of sulfur cycle metabolism in subtropical mangrove wetland ecosystems.

## CRediT authorship contribution statement

Conceptualization, C.J. and B.Y.; Methodology, S.M., B.Y. and B.L.; Software, B.Y., S.N., and J.H.; Validation, Z.Z, S.N., R.Y., and J.H.; Formal Analysis, B.Y. and C.J.; Investigation, B.Y.; Resources, C.J. and B.L.; Data Curation, J.L., and B.Y.; Writing—Original Draft Preparation, S.M.; Writing—Review and Editing, C.J.; Supervision, Q.J. and B.Y.; Project Administration, C.J.; Funding Acquisition, C.J. and B.Y.

## Acknowledgements

This research was supported by the Science and Technology Basic Resources Investigation Program of China (Grant No. 2017FY100704), the Natural Science Fund for Distinguished Young Scholars of Guangxi Zhuang Autonomous Region of China (Grant No. 2019GXNSFFA245011), the Natural Science Foundation of Guangxi Zhuang Autonomous Region of China (Grant No. 2018GXNSFAA050090, 2017GXNSFBA198101).

## References

Anantharaman, K. et al. (2018) Expanded diversity of microbial groups that shape the dissimilatory sulfur cycle. ISME J, 12, 1715–1728. DOI: 10.1038/s41396-018-0078-0.

Bender, D. et al. (2019) Impaired mitochondrial maturation of sulfite oxidase in a patient with severe sulfite oxidase deficiency. Hum Mol Genet, 28, 2885–2899. DOI: 10.1093/hmg/ddz109.

Buchfink, B. et al. (2015) Fast and sensitive protein alignment using DIAMOND. Nat Methods, 12, 59–60. DOI: 10.1038/nmeth.3176.

Bukhtiyarova, P.A. et al. (2019) Isolation, characterization, and genome insights into an anaerobic sulfidogenic Tissierella bacterium from Cu-bearing coins. Anaerobe, 56, 66–77. DOI: 10.1016/j.anaerobe.2019.02.012.

Caspi, R., et al. (2016) The MetaCyc database of metabolic pathways and enzymes and the BioCyc collection of pathway/genome databases, Nucleic Acids Res, 44(D1), D471–D480. DOI: 10.1093/nar/gkv1164.

Chiang Y.L. et al. (2009) Crystal structure of Adenylylsulfate reductase from *Desulfovibrio* gigas suggests a potential self-regulation mechanism involving the C terminus of the beta-subunit. Journal of Bacteriology, 191(24), 7597–7608. DOI: 10.1128/JB.00583-09.

Fu L. et al. (2012) CD-HIT: accelerated for clustering the next-generation sequencing data. Bioinformatics. 28(23), 3150–2. DOI: 10.1093/bioinformatics/bts565.

Ferreira, T.O. et al. (2010) Spatial patterns of soil attributes and components in a mangrove system in Southeast Brazil (Sao Paulo). J Soils Sediments, 10, 995–1006. DOI: 10.1007/s11368-010-0224-4.

Fu, L. et al. (2012) CD-HIT: accelerated for clustering the next-generation sequencing data. Bioinformatics, 28, 3150–3152. DOI: 10.1093/bioinformatics/bts565.

Fuentes-Lara, L.O. et al. (2019) From Elemental Sulfur to Hydrogen Sulfide in Agricultural Soils and Plants. Molecules, 24, 2282. DOI: 10.3390/molecules24122282.

Galperin, M.Y. et al. (2015) Expanded microbial genome coverage and improved protein family annotation in the COG database. Nucleic Acids Res, 43, D261–269. DOI: 10.1093/nar/gku1223.

George, D. and Mallery, P. (2013) IBM SPSS Statistics 21 Step by Step: A Simple Guide and Reference. Pearson, 2013, p 416.

Hausmann, B. et al. (2019) Draft Genome Sequence of *Desulfosporosinus sp*. Strain Sb-LF, Isolated from an Acidic Peatland in Germany. Microbiol Resour Announc, 8, e00428–19. DOI: 10.1128/MRA.00428-19.

Harris D. et al. (2001) Acid fumigation of soils to remove carbonates prior to total organic carbon or CARBON-13 isotopic analysis. Soil ence Society of America Journal, 65(6), 1853–1856. DOI: 10.2136/sssaj2001.1853.

Huerta-Cepas, J. et al. (2016) eggNOG 4.5: a hierarchical orthology framework with improved functional annotations for eukaryotic, prokaryotic and viral sequences. Nucleic Acids Res, 44, D286–D293. DOI: 10.1093/nar/gkv1248.

Huson D.H. et al. (2016) MEGAN Community Edition - Interactive Exploration and Analysis of Large-Scale Microbiome Sequencing Data. PLoS Comput Biol. 12(6), e1004957. DOI: 10.1371/journal.pcbi.1004957.

Hyatt D. et al. (2010) Prodigal: prokaryotic gene recognition and translation initiation site identification. BMC Bioinformatics. 11, 119. DOI: 10.1186/1471-2105-11-119.

Jochum, L.M. et al. (2018) Single-Cell Genomics Reveals a Diverse Metabolic Potential of Uncultivated Desulfatiglans-Related Deltaproteobacteria Widely Distributed in Marine Sediment. Front Microbiol, 9, 2038. DOI: 10.3389/fmicb.2018.02038.

Jorgensen, B.B. et al. (2019) The Biogeochemical Sulfur Cycle of Marine Sediments. Front Microbiol, 10, 849. DOI: 10.3389/fmicb.2019.00849.

Joshi NA, F.J. (2011) Sickle: A sliding-window, adaptive, quality-based trimming tool for FastQ files [Software]. Available at https://github.com/najoshi/sickle.

Kanehisa, M. et al. (2016) KEGG as a reference resource for gene and protein annotation. Nucleic Acids Res, 44, D457–D462. DOI: 10.1093/nar/gkv1070.

Koch, T. and Dahl, C. (2018) A novel bacterial sulfur oxidation pathway provides a new link between the cycles of organic and inorganic sulfur compounds. ISME J, 12, 2479–2491. DOI: 10.1038/s41396-018-0209-7.

Li D. et al. (2016) MEGAHIT v1.0: A fast and scalable metagenome assembler driven by advanced methodologies and community practices. Methods. 102, 3–11. DOI: 10.1016/j.ymeth.2016.02.020.

Lin, X.L. et al. (2019) Mangrove Sediment Microbiome: Adaptive Microbial Assemblages and Their Routed Biogeochemical Processes in Yunxiao Mangrove National Nature Reserve, China. Microb Ecol, 78, 57–69. DOI: 10.1007/s00248-018-1261-6.

Overbeek, R. et al. (2005) The subsystems approach to genome annotation and its use in the project to annotate 1000 genomes. Nucleic Acids Research, 33, 5691–5702. DOI: 10.1093/nar/gki866.

Peng W. et al. (2018) Metagenome complexity and template length are the main causes of bias in PCR-based bacteria community analysis. J Basic Microbiol. 58(11), 987–997. DOI: 10.1002/jobm.201800265.

Pott A.S. et al. (1998) Sirohaem sulfite reductase and other proteins encoded by genes at the dsr locus of Chromatium vinosum are involved in the oxidation of intracellular sulfur. Microbiology, 144(7), 1881–1894. DOI: 10.1099/00221287-144-7-1881.

Santos, A.A. et al. (2015) A protein trisulfide couples dissimilatory sulfate reduction to energy conservation. Science, 350, 1541–1545. DOI: 10.1126/science.aad3558.

Shi, J.C. et al. (2016) The Variation Characteristic of Sulfides and VOSc in a Source Water Reservoir and Its Control Using a Water-Lifting Aerator. Int J Environ Res Public Health, 13, 427. DOI: 10.3390/ijerph13040427.

Spring, S. et al. (2019) Sulfate-Reducing Bacteria That Produce Exopolymers Thrive in the Calcifying Zone of a Hypersaline Cyanobacterial Mat. Front Microbiol, 10, 862. DOI: 10.3389/fmicb.2019.00862

Stoffels, L. et al. (2012) Thiosulfate Reduction in Salmonella enterica Is Driven by the Proton Motive Force. J Bacteriol, 194, 475–485. DOI: 10.1128/JB.06014-11.

Tu, Q.C. et al. (2019) NCycDB: a curated integrative database for fast and accurate metagenomic profiling of nitrogen cycling genes. Bioinformatics, 35, 1040–1048. DOI: 10.1093/bioinformatics/bty741.

Varon-Lopez, M. et al. (2014) Sulphur-oxidizing and sulphate-reducing communities in Brazilian mangrove sediments. Environ Microbiol, 16, 845–855. DOI: 10.1111/1462-2920.12237.

Vavourakis, C.D. et al. (2019) Metagenomes and metatranscriptomes shed new light on the microbial-mediated sulfur cycle in a Siberian soda lake. BMC Biol, 17:69. DOI: 10.1186/s12915-019-0688-7.

Meyer B., et al. (2007) Molecular analysis of the diversity of sulfate-reducing and sulfur-oxidizing prokaryotes in the environment, using *aprA* as functional marker gene. Appl Environ Microbiol. 73(23), 7664–79. DOI: 10.1128/AEM.01272-07.

Wasmund, K. et al. (2017) The life sulfuric: microbial ecology of sulfur cycling in marine sediments. Environ Microbiol Rep, 9, 323–344. DOI: 10.1111/1758-2229.12538.

Watanabe, T. et al. (2013) Diversity of sulfur-cycle prokaryotes in freshwater lake sediments investigated using aprA as the functional marker gene. Syst Appl Microbiol, 36, 436–443. DOI: 10.1016/j.syapm.2013.04.009.

Wenk, C.B. et al. (2018) Electron carriers in microbial sulfate reduction inferred from experimental and environmental sulfur isotope fractionations. ISME J, 12, 495–507. DOI: 10.1038/ismej.2017.185.

Wilke, A. et al. (2012) The M5nr: a novel non-redundant database containing protein sequences and annotations from multiple sources and associated tools. BMC Bioinformatics 13, 141. DOI: 10.1186/1471-2105-13-141.

Yang Y. et al. (2013) Determination of sulfate in coastal salt marsh sediments with high-chloride concentration by ion chromatography: a revised method. Instrumentation ence & Technology, 41(1), 37–47. DOI: 10.1080/10739149.2012.717330.

Yang T.T. et al. (2017) Changes in microbial community composition following phytostabilization of an extremely acidic Cu mine tailings. Soil Biol Biochem, 114, 52–8. DOI: 10.1016/j.soilbio.2017.07.004.

Wu, J. et al. (2017) Oxygen Reduction Reaction Affected by Sulfate-Reducing Bacteria: Different Roles of Bacterial Cells and Metabolites. Indian J Microbiol, 57, 344–350. DOI: 10.1007/s12088-017-0667-z.

Wu, S. et al. (2019) Depth-related change of sulfate-reducing bacteria community in mangrove sediments: The influence of heavy metal contamination. Mar Pollut Bull, 140, 443–450. DOI: 10.1016/j.marpolbul.2019.01.042.

Chemistry, M. et al. (2000) A Simplifed Functional Version of the *Escherichia coli* Sulfite Reductase. J Biol Chem, 275, 37651–37656. DOI: 10.1074/jbc.M005619200.

Hou J., et al. (2020) Microbial succession during the transition from active to inactive stages of deep-sea hydrothermal vent sulfide chimneys. Microbiome. 8(1), 102. DOI: 10.1186/s40168-020-00851-8.

