## Supplementary Figure S1 for "Impacts of *Desulfobacterales* and *Chromatiales* on sulfate reduction in the subtropical mangrove ecosystem as revealed by SMDB analysis"

**Supplementary Figure S1**. Flowchart of major steps for SMDB construction. First, a core database was constructed for selected gene (sub) families by retrieving protein sequences from UniProt databases using keywords. Second, a full database was constructed by integrating target genes from databases including COG, eggNOG, KEGG, M5nr and NR. Third, the tools were developed to generate functional profiles for shotgun metagenomes.

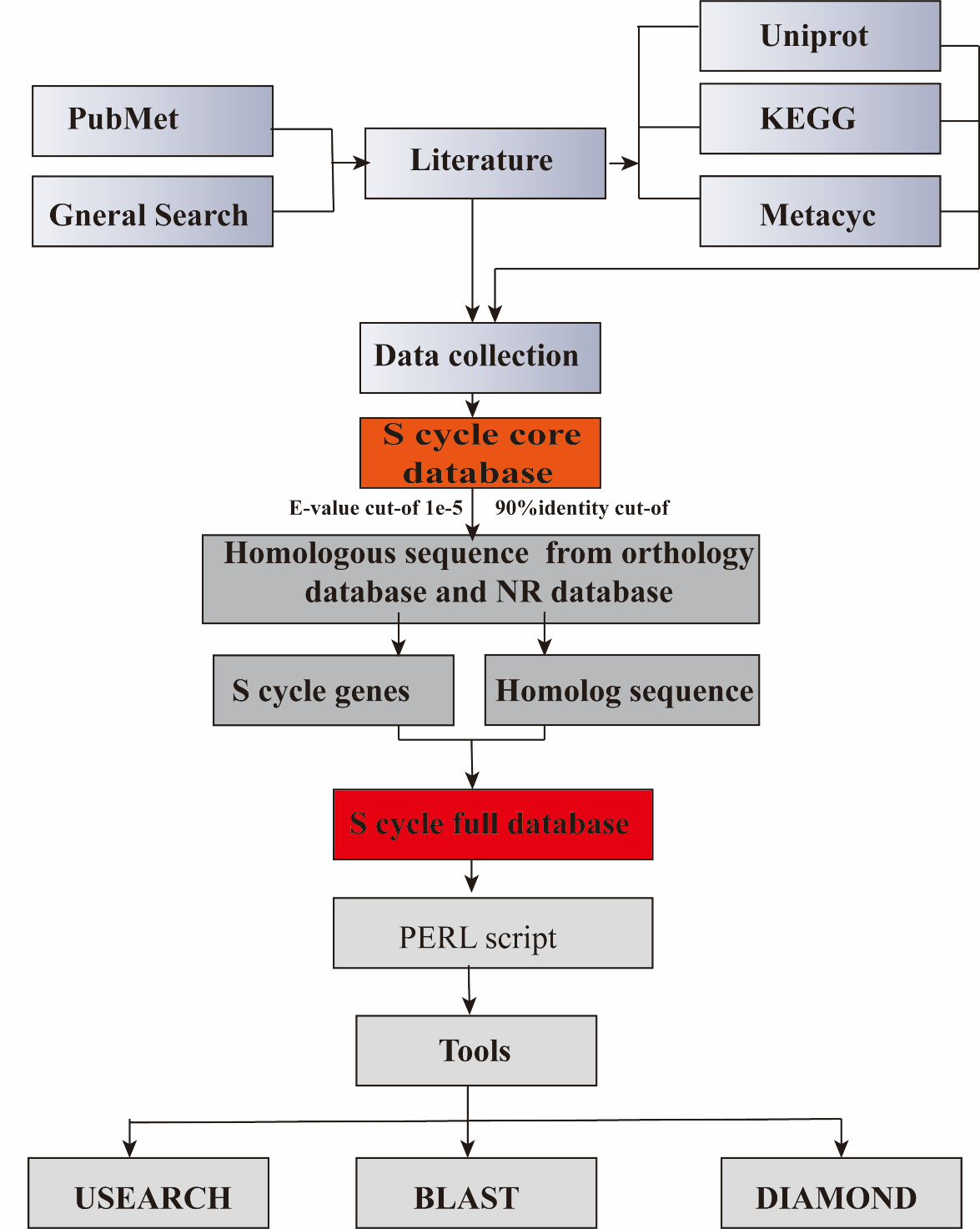
