## Supplementary Figure S2 for "Impacts of *Desulfobacterales* and *Chromatiales* on sulfate reduction in the subtropical mangrove ecosystem as revealed by SMDB analysis"

**Supplementary Figure S2**. The percentage of sequences belonging to the selected sulfur cycle gene (sub) families in public databases. Blue indicates fewer sequences of the gene (sub) family from the corresponding public database. Heatmap according to z-scores of abundant gene (sub) family.

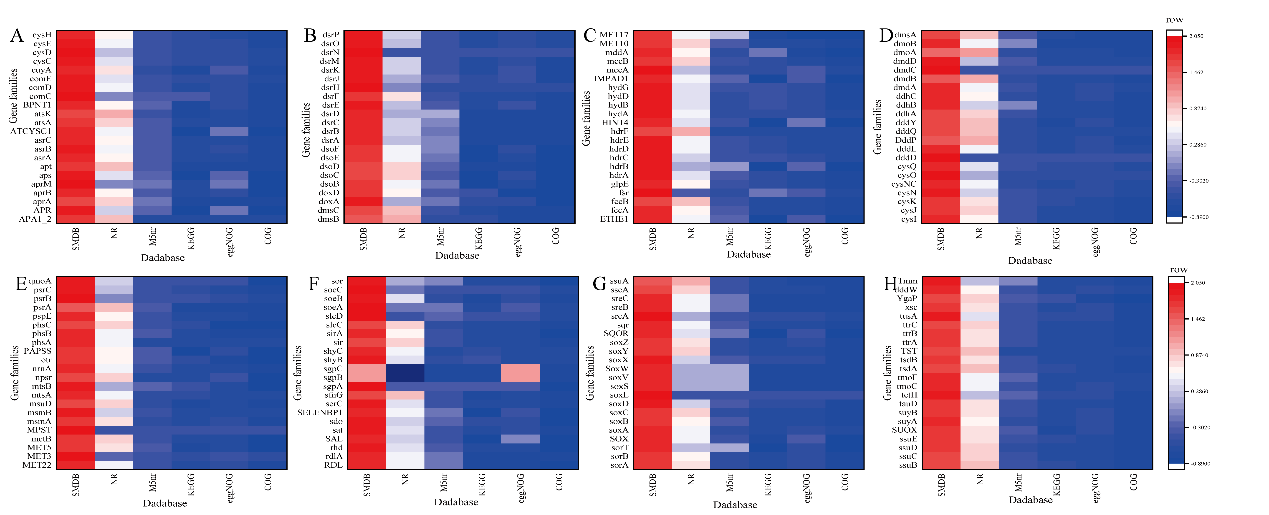
