## Supplementary Figure S3 for "Impacts of *Desulfobacterales* and *Chromatiales* on sulfate reduction in the subtropical mangrove ecosystem as revealed by SMDB analysis"

**Supplementary Figure S3**. The results of detected gene families in the mangrove wetland ecosystem in Beibu Gulf Sampling metagenome datasets by searching against SMDB, COG, eggNOG, KEGG, M5nr, and NR databases.

|  | Detected gene families |
| --- | --- |
| SMDB | 152 |
| COG | 110 |
| eggNOG | 136 |
| KEGG | 129 |
| M5nr | 147 |
| NR | 151 |
