## Supplementary Figure S4 for "Impacts of *Desulfobacterales* and *Chromatiales* on sulfate reduction in the subtropical mangrove ecosystem as revealed by SMDB analysis"

**Supplementary Figure S4**. Geographic distribution of sampling sites in the subtropical mangrove ecosystem of Beibu Gulf in China. Sediment samples were collected from National Shankou Natural Reserve of Mangrove in Beihai city (Site: 21°29′25.74″N, 109°45′49.43″E). The blue labels NMS1, NMS2, and NMS3 are the sampling sites of the nonmangrove area. The red labels RS1, RS2, and RS3 are the sampling sites of the rhizosphere area, and the yellow labels NRS1, NRS2, and NRS3 are the sampling sites of the non-rhizosphere area from near-root (within 3 cm from the rhizosphere roots). The mangrove sampling sites included rhizosphere sampling sites and non-rhizosphere sampling sites.

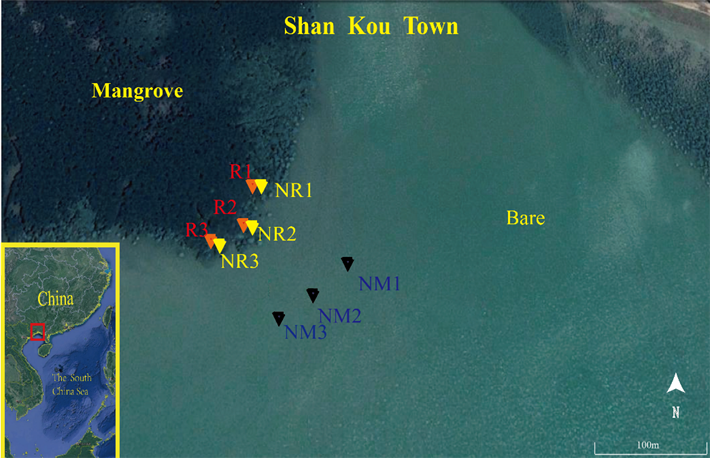
