## Supplementary Figure S5 for "Impacts of *Desulfobacterales* and *Chromatiales* on sulfate reduction in the subtropical mangrove ecosystem as revealed by SMDB analysis"

**Supplementary Figure S5**. The 20 dominant dissimilatory sulfate-reduction genes taxonomy order level are shown with their relative abundances.

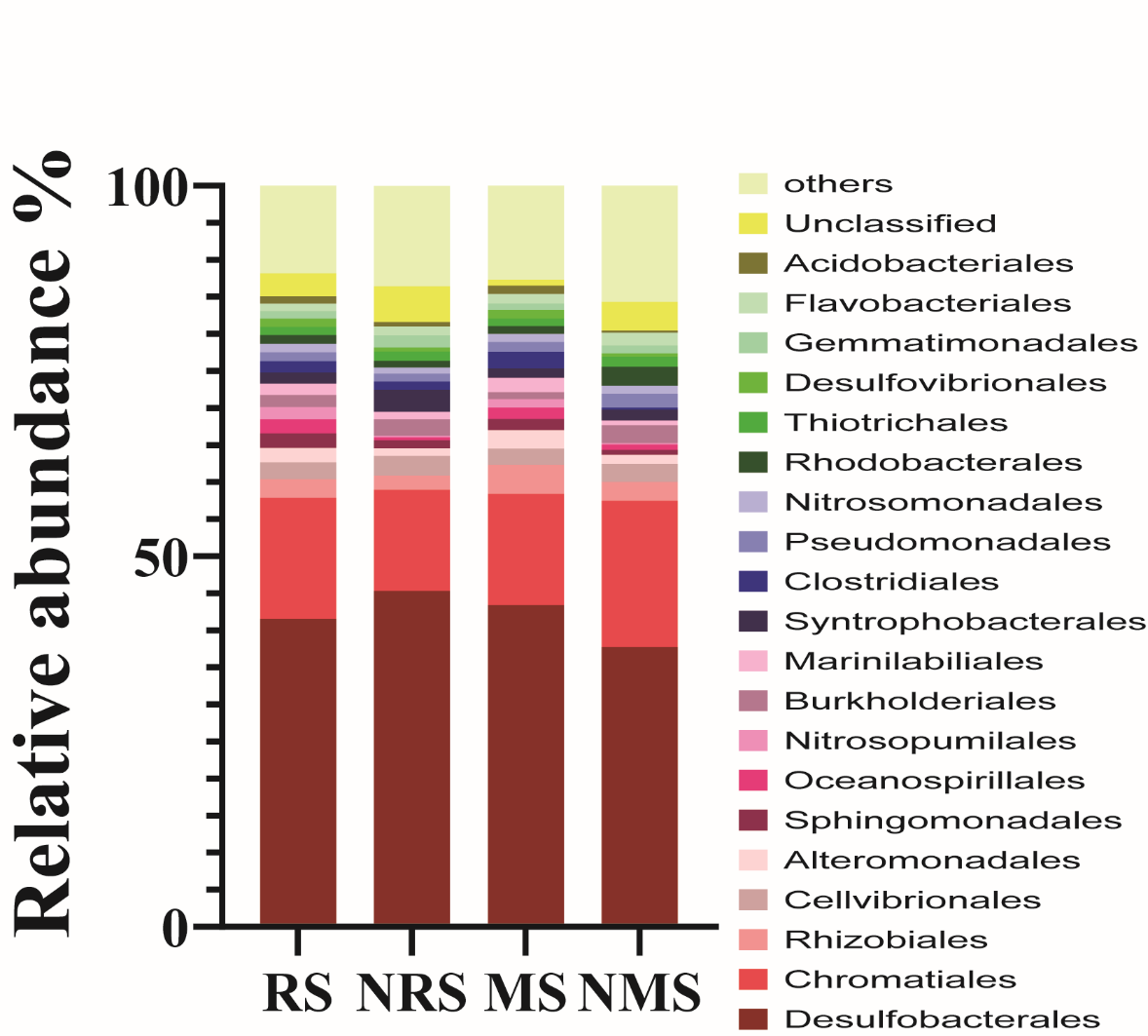
