## Supplementary Figure S6 for "Impacts of *Desulfobacterales* and *Chromatiales* on sulfate reduction in the subtropical mangrove ecosystem as revealed by SMDB analysis"

**Supplementary Figure S6**. Sediment properties. (A) TOC (mg/kg), TN (mg/kg), and TP (mg/kg). (B) Contents (mg/kg) of AS and sulfide. (C) pH, salinity, and ORP. (D) Contents (mg/g) of NH_4_^+^, NO_3_^−^ , and Fe.

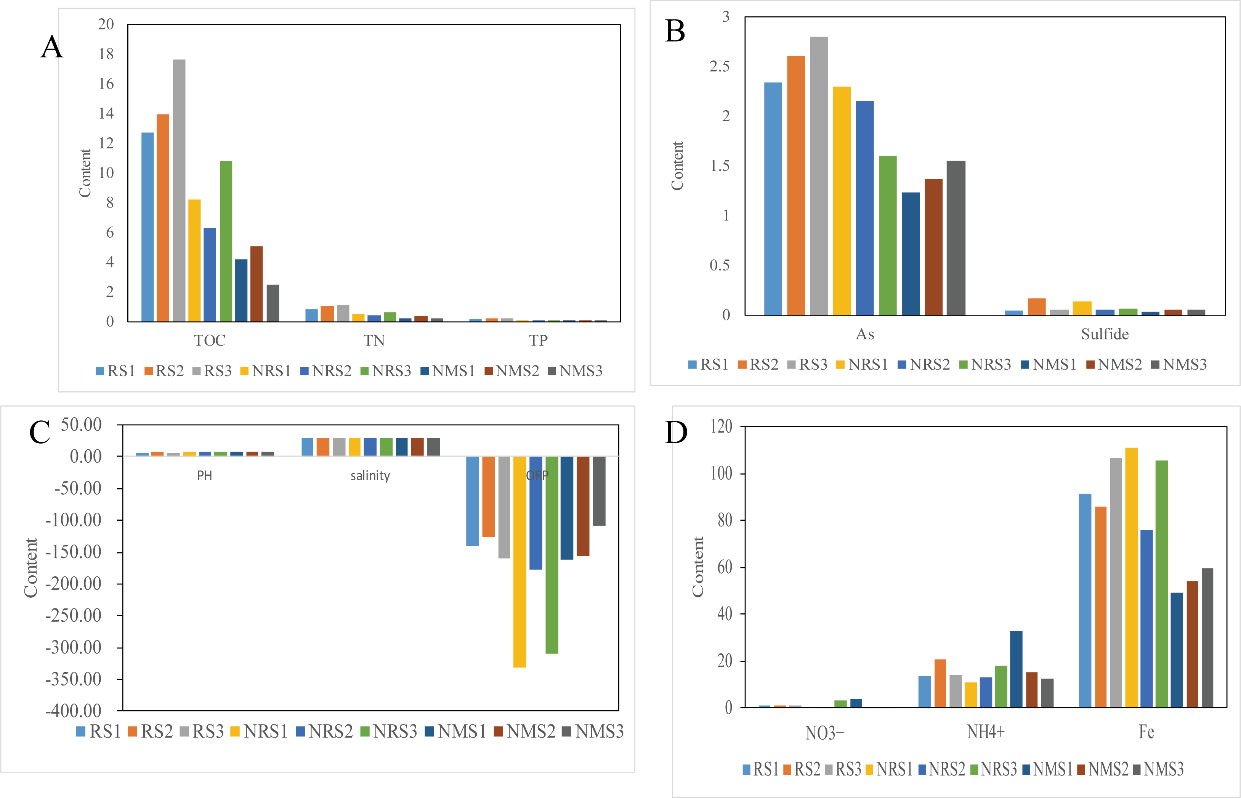
