## Supplementary Figure S7 for "Impacts of *Desulfobacterales* and *Chromatiales* on sulfate reduction in the subtropical mangrove ecosystem as revealed by SMDB analysis"

**Supplementary Figure S7**. Taxonomic classification of key functional genes retrieved from the samples. A) The key genes enriched in the MS; B) The key genes enriched in the NMS.

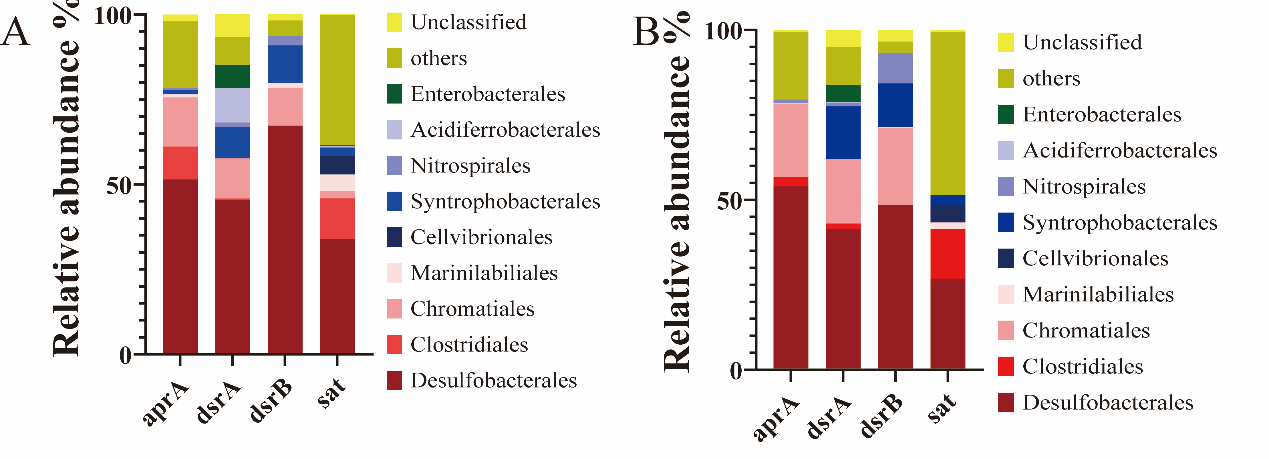
