## Supplementary Figure S8 for "Impacts of *Desulfobacterales* and *Chromatiales* on sulfate reduction in the subtropical mangrove ecosystem as revealed by SMDB analysis"

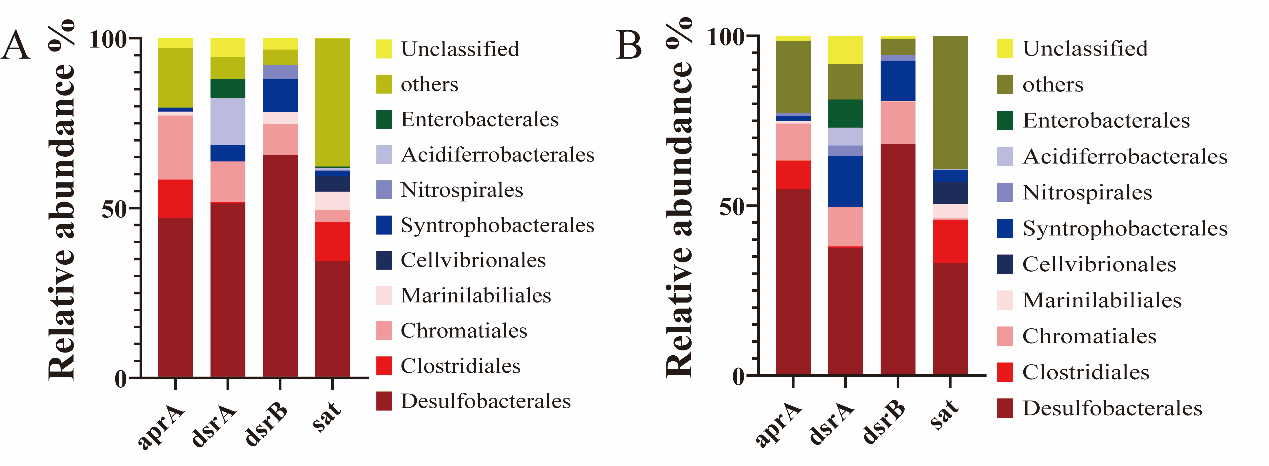
**Supplementary Figure S8**. Taxonomic classification of key functional genes retrieved from the samples. A) The key genes enriched in the RS; B) The key genes enriched in the NRS.
