## Supplementary Table S1 for "Impacts of *Desulfobacterales* and *Chromatiales* on sulfate reduction in the subtropical mangrove ecosystem as revealed by SMDB analysis"

### Supplementary Table S1. Summary of representative sequences for selected gene families in SMDB.

| gene | SMDB | NR | M5nr | KEGG | eggNOG | COG |
| --- | --- | --- | --- | --- | --- | --- |
| *APA1_2* | 9 | 6 | 0 | 0 | 0 | 0 |
| *APR* | 152 | 69 | 36 | 0 | 38 | 0 |
| *aprA* | 3767 | 2483 | 1041 | 30 | 50 | 14 |
| *aprB* | 956 | 517 | 182 | 81 | 104 | 22 |
| *aprM* | 33 | 10 | 8 | 3 | 8 | 0 |
| *aps* | 371 | 172 | 80 | 0 | 84 | 0 |
| *apt* | 6 | 4 | 1 | 0 | 0 | 0 |
| *asrA* | 2931 | 1499 | 423 | 125 | 126 | 8 |
| *asrB* | 2793 | 1359 | 378 | 138 | 157 | 18 |
| *asrC* | 2203 | 1175 | 363 | 146 | 117 | 18 |
| *ATCYSC1* | 58 | 29 | 10 | 0 | 15 | 0 |
| *atsA* | 21350 | 14005 | 3094 | 817 | 768 | 95 |
| *atsK* | 366 | 271 | 45 | 5 | 22 | 0 |
| *BPNT1* | 915 | 503 | 168 | 0 | 89 | 0 |
| *comC* | 31 | 9 | 6 | 6 | 4 | 2 |
| *comD* | 670 | 317 | 78 | 36 | 52 | 12 |
| *comE* | 511 | 231 | 47 | 31 | 35 | 12 |
| *cuyA* | 39 | 23 | 4 | 2 | 7 | 2 |
| *cysC* | 38279 | 15562 | 3098 | 1433 | 1636 | 307 |
| *cysD* | 17262 | 5633 | 1045 | 438 | 621 | 97 |
| *cysE* | 42560 | 20639 | 5322 | 2740 | 2492 | 358 |
| *cysH* | 27398 | 14130 | 2929 | 1462 | 1312 | 199 |
| *cysI* | 18365 | 9782 | 2031 | 945 | 780 | 123 |
| *cysJ* | 23711 | 15032 | 3045 | 1285 | 704 | 121 |
| *cysK* | 27175 | 17066 | 4219 | 2388 | 1781 | 279 |
| *cysN* | 21823 | 8823 | 1576 | 642 | 855 | 127 |
| *cysNC* | 2563 | 1286 | 111 | 47 | 85 | 8 |
| *cysO* | 8 | 3 | 1 | 1 | 1 | 1 |
| *cysQ* | 40999 | 18616 | 3460 | 1645 | 1826 | 274 |
| *dddD* | 6 | 1 | 1 | 1 | 1 | 0 |
| *dddL* | 317 | 156 | 15 | 3 | 11 | 1 |
| *dddP* | 674 | 511 | 45 | 7 | 38 | 3 |
| *dddQ* | 107 | 75 | 6 | 2 | 5 | 1 |
| *dddW* | 75 | 40 | 7 | 3 | 4 | 1 |
| *dddY* | 30 | 22 | 2 | 0 | 2 | 0 |
| *ddhA* | 220 | 149 | 25 | 7 | 8 | 2 |
| *ddhB* | 49 | 22 | 14 | 3 | 3 | 1 |
| *ddhC* | 21 | 11 | 2 | 1 | 2 | 0 |
| *dmdA* | 1029 | 487 | 71 | 9 | 39 | 5 |
| *dmdB* | 2799 | 2179 | 299 | 113 | 112 | 5 |
| *dmdC* | 6 | 1 | 1 | 1 | 1 | 1 |
| *dmdD* | 10 | 4 | 2 | 1 | 1 | 1 |
| *dmoA* | 540 | 461 | 53 | 6 | 17 | 0 |
| *dmoB* | 4 | 2 | 1 | 0 | 0 | 0 |
| *dmsA* | 24494 | 17993 | 3848 | 873 | 358 | 88 |
| *dmsB* | 3002 | 2293 | 368 | 266 | 60 | 12 |
| *dmsC* | 9942 | 6922 | 1470 | 576 | 147 | 34 |
| *doxA* | 8 | 3 | 2 | 1 | 1 | 0 |
| *doxD* | 383 | 227 | 72 | 65 | 9 | 0 |
| *dsoB* | 11 | 6 | 3 | 0 | 1 | 0 |
| *dsoC* | 42 | 28 | 8 | 0 | 5 | 0 |
| *dsoD* | 303 | 209 | 75 | 4 | 14 | 0 |
| *dsoE* | 11 | 6 | 3 | 0 | 1 | 0 |
| *dsoF* | 17 | 9 | 5 | 0 | 2 | 0 |
| *dsrA* | 16460 | 6390 | 4589 | 73 | 109 | 41 |
| *dsrB* | 19925 | 8582 | 4458 | 76 | 108 | 34 |
| *dsrC* | 7 | 3 | 2 | 0 | 0 | 0 |
| *dsrD* | 3 | 1 | 1 | 0 | 0 | 0 |
| *dsrE* | 40 | 17 | 8 | 4 | 7 | 3 |
| *dsrF* | 2566 | 1567 | 302 | 142 | 126 | 22 |
| *dsrH* | 7 | 2 | 1 | 1 | 1 | 1 |
| *dsrJ* | 115 | 39 | 17 | 7 | 11 | 0 |
| *dsrK* | 341 | 142 | 51 | 19 | 43 | 11 |
| *dsrM* | 187 | 75 | 23 | 5 | 14 | 2 |
| *dsrN* | 6 | 1 | 1 | 1 | 1 | 1 |
| *dsrO* | 133 | 47 | 11 | 3 | 8 | 1 |
| *dsrP* | 205 | 86 | 21 | 8 | 15 | 4 |
| *ETHE1* | 489 | 254 | 99 | 4 | 78 | 1 |
| *fccA* | 338 | 176 | 46 | 13 | 15 | 6 |
| *fccB* | 5273 | 3638 | 597 | 318 | 185 | 36 |
| *fsr* | 16 | 3 | 2 | 4 | 3 | 1 |
| *glpE* | 23382 | 12065 | 2945 | 1895 | 1221 | 154 |
| *hdrA* | 1467 | 641 | 216 | 134 | 102 | 52 |
| *hdrB* | 163 | 61 | 53 | 7 | 36 | 2 |
| *hdrC* | 1112 | 453 | 129 | 125 | 84 | 43 |
| *hdrD* | 502 | 235 | 52 | 39 | 22 | 19 |
| *hdrE* | 104 | 51 | 12 | 9 | 5 | 6 |
| *hdrF* | 4 | 3 | 0 | 0 | 0 | 0 |
| *HINT4* | 18 | 9 | 3 | 0 | 5 | 0 |
| *hydA* | 37 | 18 | 3 | 4 | 3 | 2 |
| *hydB* | 68 | 31 | 8 | 10 | 8 | 2 |
| *hydD* | 59 | 27 | 7 | 8 | 6 | 3 |
| *hydG* | 201 | 92 | 27 | 27 | 21 | 8 |
| *IMPAD1* | 563 | 277 | 114 | 0 | 86 | 0 |
| *mccA* | 18 | 7 | 2 | 2 | 3 | 1 |
| *mccB* | 1827 | 1198 | 257 | 183 | 86 | 13 |
| *mddA* | 152 | 83 | 16 | 40 | 6 | 1 |
| *MET10* | 146 | 93 | 24 | 5 | 8 | 0 |
| *MET17* | 104 | 54 | 44 | 3 | 0 | 0 |
| *MET22* | 280 | 145 | 38 | 11 | 21 | 0 |
| *MET3* | 1053 | 118 | 34 | 16 | 40 | 1 |
| *MET5* | 128 | 80 | 21 | 2 | 8 | 0 |
| *metB* | 23601 | 15207 | 3360 | 1495 | 1220 | 149 |
| *MPST* | 3 | 0 | 0 | 0 | 0 | 0 |
| *msmA* | 144 | 84 | 13 | 7 | 11 | 0 |
| *msmB* | 19 | 8 | 2 | 1 | 1 | 1 |
| *msuD* | 3117 | 2192 | 324 | 124 | 180 | 6 |
| *mtsA* | 78 | 37 | 5 | 4 | 2 | 1 |
| *mtsB* | 13 | 5 | 3 | 2 | 1 | 0 |
| *npsr* | 15 | 10 | 1 | 1 | 2 | 0 |
| *nrnA* | 18124 | 9477 | 2240 | 964 | 895 | 117 |
| *otr* | 57 | 33 | 11 | 8 | 3 | 0 |
| *PAPSS* | 1847 | 1055 | 349 | 0 | 197 | 0 |
| *phsA* | 8 | 4 | 0 | 0 | 0 | 0 |
| *phsB* | 2031 | 1021 | 343 | 153 | 72 | 24 |
| *phsC* | 1377 | 916 | 235 | 133 | 20 | 6 |
| *pspE* | 6429 | 3573 | 886 | 490 | 320 | 39 |
| *psrA* | 279 | 208 | 53 | 1 | 6 | 0 |
| *psrB* | 33 | 10 | 6 | 6 | 4 | 3 |
| *psrC* | 28 | 11 | 5 | 4 | 3 | 2 |
| *qmoA* | 7 | 3 | 1 | 1 | 1 | 0 |
| *RDL* | 42 | 22 | 10 | 2 | 2 | 0 |
| *rdlA* | 29 | 15 | 7 | 0 | 0 | 0 |
| *rhd* | 29 | 15 | 5 | 3 | 3 | 1 |
| *SAL* | 101 | 47 | 17 | 0 | 30 | 0 |
| *sat* | 48516 | 21785 | 4434 | 2118 | 2606 | 371 |
| *sdo* | 9 | 4 | 2 | 1 | 1 | 0 |
| *SELENBP1* | 429 | 223 | 108 | 0 | 65 | 0 |
| *sfnG* | 542 | 405 | 73 | 22 | 23 | 2 |
| *sgpA* | 5 | 1 | 1 | 1 | 1 | 0 |
| *sgpB* | 1 | 0 | 0 | 0 | 0 | 0 |
| *sgpC* | 1 | 0 | 0 | 0 | 0 | 0 |
| *shyB* | 28 | 12 | 1 | 2 | 1 | 0 |
| *shyC* | 81 | 38 | 2 | 3 | 1 | 2 |
| *sir* | 13452 | 9048 | 1649 | 631 | 758 | 68 |
| *sirA* | 511 | 282 | 86 | 39 | 45 | 5 |
| *slcC* | 1441 | 973 | 143 | 42 | 54 | 9 |
| *slcD* | 17 | 3 | 4 | 3 | 3 | 2 |
| *SoeA* | 19 | 5 | 5 | 2 | 4 | 2 |
| *SoeB* | 42 | 21 | 6 | 3 | 6 | 3 |
| *soeC* | 12 | 3 | 2 | 2 | 2 | 1 |
| *sor* | 73 | 26 | 21 | 5 | 6 | 1 |
| *sorA* | 1604 | 973 | 134 | 67 | 113 | 29 |
| *sorB* | 933 | 533 | 92 | 39 | 62 | 8 |
| *sorT* | 63 | 26 | 25 | 1 | 9 | 0 |
| *SOX* | 156 | 81 | 24 | 3 | 27 | 1 |
| *soxA* | 2632 | 1358 | 215 | 89 | 154 | 27 |
| *soxB* | 1784 | 1021 | 322 | 110 | 128 | 43 |
| *SoxC* | 1256 | 816 | 105 | 43 | 74 | 12 |
| *soxD* | 401 | 179 | 72 | 43 | 42 | 16 |
| *soxl* | 6 | 1 | 1 | 1 | 1 | 1 |
| *soxS* | 6 | 2 | 2 | 0 | 0 | 0 |
| *soxV* | 6 | 2 | 2 | 0 | 0 | 0 |
| *soxW* | 6 | 2 | 2 | 0 | 0 | 0 |
| *soxX* | 1922 | 860 | 314 | 177 | 200 | 58 |
| *soxY* | 2961 | 1861 | 230 | 65 | 152 | 12 |
| *soxZ* | 947 | 573 | 59 | 23 | 59 | 0 |
| *SQOR* | 361 | 178 | 101 | 1 | 63 | 0 |
| *sqr* | 1322 | 710 | 229 | 97 | 81 | 29 |
| *sreA* | 57 | 20 | 9 | 8 | 8 | 5 |
| *sreB* | 4 | 2 | 1 | 0 | 0 | 0 |
| *sreC* | 4 | 2 | 1 | 0 | 0 | 0 |
| *sseA* | 9732 | 6382 | 1526 | 791 | 471 | 85 |
| *ssuA* | 7824 | 5984 | 977 | 539 | 159 | 42 |
| *ssuB* | 5615 | 4115 | 736 | 430 | 193 | 30 |
| *ssuC* | 20069 | 13582 | 2942 | 1445 | 792 | 103 |
| *ssuD* | 62904 | 39079 | 5765 | 2266 | 2976 | 272 |
| *ssuE* | 9731 | 5639 | 1159 | 644 | 484 | 53 |
| *SUOX* | 971 | 573 | 168 | 15 | 100 | 1 |
| *suyA* | 386 | 204 | 40 | 20 | 35 | 7 |
| *suyB* | 1196 | 749 | 155 | 51 | 88 | 15 |
| *tauD* | 36860 | 21645 | 4311 | 1992 | 1798 | 192 |
| *tetH* | 30 | 12 | 7 | 3 | 3 | 1 |
| *Tmm* | 81 | 29 | 21 | 3 | 3 | 1 |
| *tmoC* | 98 | 48 | 10 | 2 | 7 | 1 |
| *tmoF* | 6 | 3 | 1 | 0 | 0 | 0 |
| *tsdA* | 547 | 384 | 67 | 41 | 20 | 6 |
| *tsdB* | 127 | 76 | 18 | 10 | 11 | 3 |
| *TST* | 19 | 14 | 1 | 1 | 1 | 1 |
| *ttrA* | 5050 | 3132 | 584 | 150 | 131 | 20 |
| *ttrB* | 3322 | 2054 | 377 | 147 | 140 | 32 |
| *ttrC* | 2319 | 1545 | 287 | 99 | 45 | 7 |
| *tusA* | 27040 | 12360 | 2816 | 1967 | 1655 | 250 |
| *xsc* | 4244 | 2423 | 388 | 128 | 233 | 36 |
| *ygaP* | 549 | 362 | 94 | 78 | 11 | 0 |
