## Supplementary Table S2 for "Impacts of *Desulfobacterales* and *Chromatiales* on sulfate reduction in the subtropical mangrove ecosystem as revealed by SMDB analysis"

### Supplementary Table S2. Functional gene abundances.

| Gene | RS1 | RS2 | RS3 | NRS1 | NRS2 | NRS3 | NMS1 | NMS2 | NMS3 |
| --- | --- | --- | --- | --- | --- | --- | --- | --- | --- |
| *ssuB* | 13063.18 | 12076.69 | 12465.27 | 13420.74 | 14340.13 | 14241.59 | 13626.71 | 14014.67 | 14153.28 |
| *hdrA* | 9296.63 | 6430.16 | 7483.20 | 10067.70 | 10136.89 | 10055.02 | 7450.07 | 8143.39 | 8859.13 |
| *atsA* | 5258.04 | 5077.12 | 4154.05 | 4772.87 | 4607.08 | 5284.68 | 5202.72 | 4722.90 | 4674.07 |
| *dmdB* | 3684.91 | 3260.91 | 3345.63 | 3698.17 | 4105.48 | 4236.99 | 3878.75 | 3963.07 | 4100.45 |
| *dmsA* | 2876.96 | 2386.96 | 2986.59 | 2677.03 | 2632.53 | 2982.90 | 2328.39 | 2314.50 | 2649.85 |
| *cysC* | 2387.02 | 2321.65 | 2102.74 | 2378.26 | 2384.64 | 2425.62 | 2329.94 | 2461.90 | 2378.28 |
| *hdrD* | 1932.85 | 1342.28 | 1715.20 | 1950.09 | 1954.98 | 1949.67 | 1357.47 | 1413.98 | 1665.73 |
| *tusA* | 1902.31 | 1793.84 | 1770.72 | 1815.25 | 1763.83 | 1988.61 | 1754.32 | 1841.03 | 1773.16 |
| *cysE* | 1842.07 | 1918.55 | 1862.49 | 1871.16 | 1915.63 | 1908.43 | 1852.13 | 1956.58 | 1860.37 |
| *ttrB* | 1415.79 | 1398.95 | 1552.77 | 1375.81 | 1433.07 | 1508.67 | 1145.23 | 1213.50 | 1307.00 |
| *cysK* | 1297.39 | 1338.91 | 1181.08 | 1148.58 | 1320.67 | 1392.06 | 1562.54 | 1589.62 | 1320.42 |
| *slcC* | 1254.05 | 1157.89 | 1056.36 | 1286.80 | 1262.99 | 1308.38 | 1436.12 | 1388.17 | 1262.03 |
| *xsc* | 1253.77 | 1195.00 | 1176.94 | 1263.65 | 1250.10 | 1277.79 | 1247.24 | 1428.72 | 1159.35 |
| *sat* | 1221.65 | 1185.42 | 1302.43 | 1140.46 | 1121.97 | 1279.84 | 1369.75 | 1288.09 | 1303.29 |
| *cysN* | 1166.10 | 1306.24 | 1104.03 | 1222.93 | 1349.45 | 1225.31 | 1372.96 | 1491.66 | 1302.46 |
| *cysQ* | 1128.33 | 1180.15 | 1019.25 | 997.92 | 1036.19 | 1200.05 | 1095.65 | 1059.73 | 1068.27 |
| *glpE* | 973.60 | 963.74 | 824.45 | 978.54 | 920.98 | 983.41 | 951.43 | 956.04 | 886.82 |
| *ssuD* | 965.61 | 1050.51 | 847.14 | 846.68 | 740.63 | 1018.34 | 1130.60 | 1152.44 | 839.07 |
| *cysJ* | 802.26 | 802.75 | 831.86 | 742.84 | 805.82 | 794.77 | 783.59 | 842.88 | 818.14 |
| *tauD* | 796.79 | 837.99 | 799.26 | 692.15 | 796.44 | 841.20 | 881.77 | 837.33 | 843.33 |
| *nrnA* | 781.83 | 610.89 | 799.94 | 820.70 | 956.30 | 801.13 | 628.13 | 695.50 | 836.54 |
| *dsrK* | 774.30 | 708.07 | 616.90 | 809.42 | 845.70 | 942.23 | 573.50 | 754.62 | 767.06 |
| *metB* | 747.04 | 659.75 | 613.80 | 714.15 | 833.35 | 695.00 | 1030.50 | 947.59 | 802.95 |
| *aprA* | 722.93 | 600.10 | 540.46 | 737.23 | 583.28 | 743.12 | 437.70 | 476.07 | 595.25 |
| *ddhA* | 670.98 | 506.85 | 742.03 | 710.56 | 724.91 | 754.57 | 513.70 | 593.00 | 609.93 |
| *pspE* | 597.58 | 625.72 | 446.81 | 516.48 | 548.51 | 524.83 | 468.20 | 488.23 | 604.07 |
| *dmdC* | 543.99 | 432.74 | 450.12 | 424.09 | 557.01 | 499.09 | 557.84 | 524.47 | 530.16 |
| *dmdA* | 535.69 | 505.59 | 425.93 | 462.19 | 627.46 | 536.04 | 886.58 | 803.75 | 668.32 |
| *cysH* | 530.74 | 429.74 | 504.71 | 477.38 | 539.81 | 521.36 | 507.08 | 463.13 | 443.30 |
| *hdrC* | 514.34 | 369.63 | 386.27 | 633.34 | 579.95 | 476.12 | 360.05 | 419.71 | 475.83 |
| *dsrP* | 448.29 | 367.68 | 526.30 | 473.36 | 428.09 | 419.95 | 266.21 | 338.10 | 372.60 |
| *sir* | 421.90 | 588.58 | 392.43 | 357.97 | 373.20 | 411.36 | 381.04 | 390.37 | 423.38 |
| *fccB* | 400.68 | 458.92 | 398.21 | 303.43 | 358.44 | 416.68 | 370.11 | 354.17 | 332.34 |
| *asrB* | 390.76 | 323.98 | 373.38 | 512.60 | 474.48 | 469.77 | 361.94 | 337.26 | 386.63 |
| *otr* | 387.53 | 282.05 | 233.13 | 354.24 | 371.34 | 442.30 | 305.91 | 352.62 | 367.31 |
| *msmA* | 385.34 | 416.92 | 319.81 | 274.15 | 261.10 | 301.52 | 426.14 | 358.34 | 309.67 |
| *soxB* | 381.57 | 509.57 | 374.19 | 321.80 | 285.65 | 345.65 | 400.48 | 405.12 | 387.75 |
| *psrA* | 366.84 | 275.21 | 386.80 | 426.07 | 382.77 | 376.53 | 274.39 | 327.96 | 321.33 |
| *dsrB* | 362.75 | 254.47 | 276.02 | 372.36 | 359.21 | 367.98 | 202.01 | 265.93 | 331.01 |
| *dmdD* | 356.56 | 238.42 | 224.75 | 410.30 | 464.24 | 379.21 | 378.55 | 406.95 | 414.61 |
| *ssuC* | 350.29 | 413.97 | 427.90 | 290.24 | 359.36 | 385.93 | 422.94 | 458.47 | 398.09 |
| *SUOX* | 314.18 | 314.19 | 267.07 | 419.92 | 287.81 | 276.19 | 329.20 | 249.23 | 317.24 |
| *dsrF* | 313.35 | 446.91 | 258.78 | 285.03 | 228.77 | 296.94 | 344.99 | 328.43 | 319.36 |
| *ssuA* | 312.98 | 334.83 | 364.51 | 349.98 | 321.84 | 330.15 | 334.90 | 344.93 | 348.01 |
| *sseA* | 299.09 | 433.70 | 296.41 | 211.87 | 291.57 | 263.71 | 356.45 | 303.53 | 261.23 |
| *dsrM* | 293.80 | 240.93 | 240.35 | 286.19 | 226.13 | 260.56 | 171.73 | 151.50 | 189.07 |
| *asrC* | 291.07 | 193.71 | 391.66 | 334.32 | 343.92 | 340.15 | 173.52 | 227.78 | 308.13 |
| *MET3* | 287.25 | 316.40 | 196.65 | 288.70 | 270.44 | 304.70 | 302.71 | 320.92 | 281.24 |
| *dsrA* | 285.79 | 291.53 | 313.44 | 304.72 | 242.39 | 307.60 | 203.73 | 244.25 | 258.19 |
| *hdrE* | 254.30 | 99.61 | 168.96 | 305.08 | 278.47 | 248.63 | 155.53 | 157.86 | 234.37 |
| *ETHE1* | 230.98 | 274.49 | 200.26 | 194.76 | 211.52 | 256.70 | 213.49 | 228.21 | 223.16 |
| *sqr* | 230.02 | 184.73 | 131.45 | 196.43 | 171.66 | 187.36 | 227.76 | 228.81 | 205.87 |
| *dsrN* | 226.69 | 191.23 | 304.58 | 287.16 | 297.03 | 226.27 | 184.37 | 211.27 | 293.07 |
| *sorA* | 214.20 | 176.83 | 143.40 | 175.01 | 236.07 | 235.99 | 284.99 | 277.67 | 257.49 |
| *asrA* | 208.76 | 144.89 | 162.31 | 232.07 | 176.79 | 159.12 | 130.46 | 181.14 | 164.60 |
| *dddD* | 198.50 | 190.67 | 167.86 | 236.52 | 219.00 | 225.97 | 202.46 | 196.29 | 165.34 |
| *MET10* | 195.03 | 194.54 | 165.19 | 161.44 | 144.56 | 167.12 | 145.32 | 170.01 | 156.72 |
| *cysI* | 191.07 | 148.69 | 152.34 | 123.31 | 155.55 | 147.63 | 179.66 | 145.82 | 158.39 |
| *fccA* | 186.13 | 454.07 | 218.92 | 129.96 | 144.30 | 161.27 | 237.92 | 201.90 | 231.41 |
| *mtsA* | 185.64 | 132.08 | 107.51 | 148.83 | 166.68 | 175.87 | 169.86 | 141.74 | 205.42 |
| *hdrB* | 182.37 | 161.43 | 134.99 | 138.21 | 130.24 | 153.37 | 227.38 | 193.25 | 209.72 |
| *cysNC* | 181.77 | 330.38 | 162.21 | 160.30 | 204.10 | 208.27 | 271.07 | 258.53 | 222.76 |
| *ttrA* | 178.79 | 123.58 | 131.96 | 189.50 | 207.93 | 176.95 | 256.09 | 220.22 | 201.31 |
| *mtsB* | 173.87 | 144.13 | 224.14 | 206.21 | 201.41 | 274.83 | 123.15 | 159.09 | 173.75 |
| *cuyA* | 164.15 | 203.07 | 92.64 | 133.69 | 189.16 | 162.83 | 252.26 | 273.43 | 223.04 |
| *comE* | 160.21 | 183.63 | 195.43 | 160.07 | 174.76 | 175.08 | 270.35 | 238.57 | 199.25 |
| *hydA* | 146.08 | 110.38 | 112.05 | 193.37 | 157.30 | 195.59 | 123.30 | 133.11 | 172.33 |
| *npsr* | 144.52 | 164.93 | 162.11 | 165.35 | 150.09 | 188.84 | 138.50 | 134.93 | 124.18 |
| *tsdB* | 144.10 | 220.79 | 135.96 | 89.62 | 93.25 | 112.26 | 135.64 | 116.22 | 167.33 |
| *phsB* | 142.79 | 111.70 | 112.37 | 182.76 | 193.88 | 147.15 | 99.89 | 87.66 | 98.60 |
| *dddP* | 136.56 | 122.11 | 194.17 | 194.60 | 152.71 | 188.52 | 179.97 | 150.85 | 175.17 |
| *phsA* | 136.25 | 71.81 | 79.09 | 161.93 | 205.39 | 208.91 | 185.60 | 220.75 | 225.10 |
| *cysD* | 133.96 | 185.17 | 126.32 | 152.22 | 153.92 | 136.44 | 208.69 | 205.09 | 175.84 |
| *aprB* | 133.65 | 170.23 | 104.25 | 151.58 | 156.94 | 146.11 | 117.16 | 73.23 | 168.78 |
| *dsrO* | 132.90 | 146.85 | 134.83 | 138.42 | 98.31 | 153.04 | 97.05 | 123.27 | 137.41 |
| *dsoF* | 124.97 | 114.51 | 85.02 | 79.43 | 80.85 | 105.07 | 75.21 | 98.10 | 87.90 |
| *dmsB* | 124.80 | 142.83 | 127.11 | 119.74 | 183.34 | 157.34 | 102.14 | 145.29 | 153.66 |
| *MET5* | 122.56 | 155.75 | 139.11 | 152.85 | 113.40 | 112.68 | 187.76 | 153.14 | 195.94 |
| *hydD* | 118.71 | 112.64 | 68.45 | 87.91 | 91.61 | 98.46 | 71.96 | 59.02 | 47.10 |
| *dsrC* | 111.58 | 86.32 | 111.19 | 105.34 | 78.18 | 134.71 | 93.25 | 72.74 | 86.25 |
| *mccB* | 111.19 | 127.99 | 86.13 | 124.94 | 153.14 | 88.14 | 140.82 | 152.67 | 146.61 |
| *MET17* | 110.93 | 100.28 | 101.86 | 126.35 | 153.23 | 141.32 | 132.74 | 91.19 | 92.88 |
| *suyB* | 109.44 | 99.17 | 102.37 | 124.40 | 152.73 | 139.52 | 181.18 | 146.92 | 164.44 |
| *soxY* | 104.26 | 138.76 | 84.32 | 108.96 | 78.59 | 94.57 | 135.81 | 156.76 | 128.85 |
| *phsC* | 103.56 | 95.31 | 63.87 | 82.19 | 84.49 | 102.17 | 84.89 | 82.12 | 68.94 |
| *soxA* | 96.85 | 154.81 | 92.42 | 113.22 | 115.16 | 146.65 | 151.86 | 141.05 | 132.13 |
| *shyC* | 91.73 | 83.35 | 53.29 | 56.40 | 62.31 | 70.82 | 49.13 | 57.76 | 51.89 |
| *aprM* | 91.18 | 138.60 | 45.36 | 48.10 | 70.42 | 45.28 | 75.97 | 69.40 | 68.51 |
| *mddA* | 81.16 | 80.97 | 58.49 | 90.01 | 92.62 | 83.80 | 88.80 | 106.80 | 117.81 |
| *sorB* | 77.45 | 153.46 | 92.91 | 58.53 | 78.15 | 58.18 | 165.92 | 111.82 | 103.60 |
| *dsrJ* | 75.44 | 108.19 | 120.37 | 89.99 | 113.16 | 90.57 | 63.88 | 81.42 | 97.84 |
| *soxX* | 75.08 | 71.82 | 39.30 | 91.34 | 45.70 | 68.45 | 83.87 | 74.61 | 90.08 |
| *soxD* | 74.82 | 94.77 | 78.58 | 75.74 | 81.68 | 75.55 | 116.79 | 118.55 | 57.80 |
| *sdo* | 74.41 | 125.96 | 103.32 | 121.49 | 117.57 | 119.01 | 174.36 | 129.40 | 198.65 |
| *psrC* | 68.01 | 25.27 | 41.18 | 33.26 | 21.85 | 61.57 | 43.52 | 55.05 | 62.94 |
| *apt* | 67.39 | 121.13 | 60.40 | 42.95 | 63.74 | 77.34 | 66.09 | 49.72 | 86.18 |
| *soxV* | 64.83 | 77.82 | 99.09 | 60.76 | 48.68 | 41.70 | 90.07 | 81.74 | 50.91 |
| *ssuE* | 62.03 | 99.99 | 108.39 | 101.55 | 144.25 | 80.10 | 67.43 | 118.44 | 60.01 |
| *ttrC* | 58.88 | 65.75 | 78.13 | 91.55 | 80.42 | 73.44 | 86.37 | 108.43 | 83.22 |
| *soxl* | 55.06 | 81.97 | 24.69 | 63.54 | 51.87 | 61.78 | 48.56 | 67.85 | 59.22 |
| *suyA* | 52.32 | 20.23 | 4.39 | 36.58 | 41.37 | 51.76 | 47.86 | 47.55 | 26.08 |
| *soxZ* | 51.24 | 102.52 | 65.46 | 55.18 | 91.98 | 55.65 | 89.09 | 60.14 | 75.27 |
| *dmsC* | 50.96 | 57.64 | 28.00 | 51.65 | 76.64 | 59.50 | 58.13 | 48.51 | 73.07 |
| *SoeA* | 48.96 | 84.16 | 27.49 | 22.48 | 46.89 | 47.66 | 28.22 | 52.74 | 48.55 |
| *comC* | 48.38 | 26.20 | 42.56 | 73.45 | 35.60 | 56.08 | 33.29 | 53.58 | 28.04 |
| *MET22* | 44.58 | 96.17 | 37.28 | 46.32 | 29.29 | 44.92 | 43.42 | 30.61 | 50.65 |
| *slcD* | 37.34 | 39.58 | 42.32 | 44.75 | 33.07 | 28.44 | 75.77 | 57.41 | 48.37 |
| *msuD* | 35.87 | 35.38 | 16.70 | 26.28 | 51.75 | 28.06 | 42.74 | 29.50 | 29.61 |
| *tmoC* | 34.57 | 17.25 | 5.63 | 9.38 | 45.77 | 7.69 | 8.74 | 15.25 | 41.60 |
| *comD* | 34.28 | 22.88 | 16.53 | 37.90 | 44.57 | 40.72 | 39.54 | 62.48 | 25.30 |
| *sreA* | 33.19 | 6.73 | 51.47 | 31.32 | 16.04 | 23.14 | 15.12 | 14.98 | 31.60 |
| *SoxC* | 32.27 | 57.85 | 37.90 | 44.27 | 64.80 | 50.44 | 68.15 | 70.87 | 51.45 |
| *soeC* | 30.81 | 38.05 | 17.17 | 8.55 | 18.54 | 29.90 | 19.90 | 16.05 | 12.29 |
| *SoeB* | 29.66 | 3.49 | 6.90 | 1.84 | 16.77 | 8.29 | 4.20 | 3.86 | 9.47 |
| *dsrE* | 27.86 | 29.52 | 33.40 | 38.53 | 18.92 | 22.78 | 25.82 | 42.13 | 19.78 |
| *psrB* | 24.02 | 18.72 | 20.35 | 30.35 | 22.95 | 12.53 | 29.62 | 25.79 | 22.10 |
| *hydG* | 23.67 | 43.90 | 23.81 | 19.32 | 11.89 | 49.20 | 20.80 | 45.25 | 42.61 |
| *SELENBP1* | 19.55 | 11.70 | 15.59 | 6.72 | 8.81 | 17.63 | 22.12 | 38.25 | 11.35 |
| *Tmm* | 18.76 | 13.41 | 5.04 | 7.00 | 14.83 | 13.94 | 28.38 | 18.60 | 29.12 |
| *doxD* | 17.87 | 4.46 | 6.98 | 18.96 | 16.20 | 20.89 | 25.13 | 19.49 | 21.91 |
| *rhd* | 14.54 | 0.00 | 4.77 | 15.72 | 0.00 | 15.79 | 2.13 | 6.58 | 18.15 |
| *soxW* | 13.88 | 2.72 | 7.58 | 15.98 | 20.49 | 22.51 | 16.79 | 14.79 | 27.73 |
| *fsr* | 13.86 | 5.04 | 6.90 | 13.60 | 8.85 | 27.28 | 2.96 | 4.15 | 15.94 |
| *dsrH* | 13.22 | 36.24 | 28.33 | 10.40 | 10.73 | 19.56 | 37.33 | 12.42 | 7.12 |
| *dddQ* | 9.72 | 4.41 | 0.00 | 5.45 | 5.78 | 7.62 | 7.04 | 1.22 | 6.41 |
| *HINT4* | 9.60 | 20.09 | 14.98 | 9.46 | 5.44 | 29.13 | 2.20 | 10.66 | 6.36 |
| *dsoD* | 9.44 | 0.60 | 7.01 | 3.51 | 4.67 | 8.14 | 4.57 | 6.91 | 14.49 |
| *tmoF* | 9.33 | 21.89 | 11.85 | 20.35 | 14.48 | 11.55 | 20.37 | 26.07 | 47.37 |
| *SOX* | 8.60 | 0.00 | 0.00 | 0.00 | 9.68 | 5.35 | 0.00 | 3.89 | 11.96 |
| *tsdA* | 8.24 | 33.85 | 20.09 | 22.66 | 2.20 | 17.69 | 28.10 | 24.60 | 21.95 |
| *dmoB* | 7.53 | 27.62 | 4.70 | 28.85 | 12.80 | 10.32 | 19.53 | 6.62 | 14.88 |
| *dmoA* | 6.86 | 0.00 | 0.00 | 0.00 | 0.00 | 0.00 | 0.00 | 4.27 | 0.00 |
| *ddhB* | 6.86 | 13.72 | 3.13 | 12.14 | 19.51 | 24.56 | 28.82 | 28.26 | 29.11 |
| *sirA* | 6.83 | 9.51 | 7.85 | 4.79 | 11.17 | 3.39 | 12.10 | 9.19 | 7.62 |
| *dddL* | 6.12 | 1.75 | 9.51 | 0.00 | 2.24 | 0.00 | 12.45 | 10.99 | 0.00 |
| *sor* | 3.36 | 0.00 | 0.00 | 3.90 | 6.81 | 4.18 | 11.92 | 4.06 | 7.57 |
| *SAL* | 3.18 | 0.00 | 0.00 | 4.69 | 0.00 | 0.00 | 0.00 | 0.00 | 0.00 |
| *shyB* | 3.15 | 1.20 | 1.57 | 6.58 | 3.85 | 1.62 | 0.00 | 0.98 | 2.82 |
| *ygaP* | 3.13 | 12.99 | 1.39 | 0.00 | 7.29 | 0.00 | 2.56 | 0.00 | 0.00 |
| *SQOR* | 2.39 | 0.91 | 2.82 | 0.78 | 0.00 | 0.73 | 1.48 | 0.00 | 2.14 |
| *dddW* | 2.05 | 9.20 | 0.00 | 5.90 | 8.96 | 29.88 | 0.00 | 1.63 | 0.00 |
| *qmoA* | 1.24 | 0.00 | 1.96 | 1.22 | 3.02 | 4.04 | 0.00 | 2.32 | 0.00 |
| *ddhC* | 1.07 | 3.66 | 0.95 | 0.00 | 1.24 | 0.00 | 1.42 | 4.29 | 0.00 |
| *soxS* | 0.00 | 0.00 | 0.00 | 10.90 | 4.49 | 1.69 | 1.72 | 0.00 | 0.00 |
| *dsoB* | 0.00 | 5.07 | 2.25 | 4.15 | 2.91 | 5.60 | 5.50 | 4.93 | 0.76 |
| *hdrF* | 0.00 | 0.00 | 6.79 | 1.88 | 0.00 | 2.89 | 8.37 | 0.00 | 2.01 |
| *TST* | 0.00 | 0.00 | 0.00 | 0.00 | 5.63 | 0.00 | 0.00 | 8.83 | 5.09 |
| *MPST* | 0.00 | 2.14 | 1.66 | 0.00 | 0.00 | 6.86 | 0.00 | 0.00 | 3.36 |
| *dsrD* | 0.00 | 0.00 | 0.00 | 19.08 | 0.00 | 0.00 | 0.00 | 0.00 | 0.00 |
| *APA1_2* | 0.00 | 0.00 | 1.70 | 0.00 | 4.75 | 0.00 | 8.63 | 8.16 | 2.10 |
| *sorT* | 0.00 | 0.00 | 1.57 | 0.00 | 0.00 | 1.46 | 5.96 | 1.65 | 4.75 |
