## Supplementary Table S3 for "Impacts of *Desulfobacterales* and *Chromatiales* on sulfate reduction in the subtropical mangrove ecosystem as revealed by SMDB analysis"

### Supplementary Table S3. Impact of sediment properties on dissimilating sulfate reduction gene families.

| Gene | PH | salinity | ORP | AS | NO_3_^-^ | NH_4_^+^ | Fe | TOC | TN | TP | Sulfide |
| --- | --- | --- | --- | --- | --- | --- | --- | --- | --- | --- | --- |
| *sat* | 0.15 | 0.13 | 0.30 | -0.56 | 0.55 | 0.56 | -0.45 | -0.16 | -0.24 | -0.01 | -0.59 |
| *cysN* | 0.76* | 0.78* | 0.24 | -0.68* | 0.06 | 0.30 | -0.82** | -0.74* | -0.67* | -0.44 | -0.09 |
| *cysQ* | 0.17 | -0.25 | 0.05 | -0.14 | 0.48 | 0.42 | 0.03 | 0.22 | 0.23 | 0.12 | 0.13 |
| *cysJ* | -0.24 | 0.21 | 0.68* | -0.07 | -0.28 | -0.10 | -0.37 | 0.09 | 0.14 | 0.47 | -0.42 |
| *nrnA* | -0.47 | -0.07 | -0.27 | 0.18 | -0.42 | -0.72* | 0.32 | -0.10 | -0.12 | -0.25 | -0.26 |
| *dsrK* | 0.00 | -0.20 | -0.55 | -0.08 | -0.18 | -0.55 | 0.36 | -0.11 | -0.13 | -0.34 | 0.10 |
| *aprA* | -0.24 | -0.71* | -0.59 | 0.36 | -0.17 | -0.53 | 0.75* | 0.32 | 0.29 | -0.06 | 0.34 |
| *cysH* | -0.56 | 0.24 | -0.28 | 0.03 | 0.33 | 0.02 | 0.24 | 0.15 | 0.03 | -0.25 | -0.58 |
| *dsrP* | -0.83** | -0.58 | -0.34 | .767* | -0.48 | -0.72* | 0.86** | 0.66 | 0.63 | 0.45 | 0.16 |
| *sir* | 0.15 | -0.37 | 0.40 | 0.34 | -0.08 | 0.20 | 0.03 | 0.38 | 0.50 | 0.57 | 0.62 |
| *asrB* | -0.18 | -0.14 | -0.81** | 0.07 | -0.02 | -0.44 | 0.53 | -0.11 | -0.19 | -0.51 | 0.06 |
| *dsrB* | -0.29 | -0.42 | -0.51 | 0.22 | -0.39 | -0.76* | 0.56 | 0.04 | 0.02 | -0.25 | 0.05 |
| *dsrM* | -0.52 | -0.69* | -0.48 | 0.68* | -0.15 | -0.40 | 0.88** | 0.63 | 0.60 | 0.21 | 0.39 |
| *asrC* | -0.74* | -0.41 | -0.39 | 0.43 | -0.36 | -0.72* | 0.67 | 0.33 | 0.26 | 0.13 | -0.15 |
| *MET3* | 0.84** | 0.25 | -0.09 | -0.56 | 0.24 | 0.31 | -0.41 | -0.49 | -0.44 | -0.47 | 0.24 |
| *dsrA* | -0.56 | -0.85** | -0.43 | 0.69* | -0.30 | -0.54 | 0.92** | 0.76* | 0.74* | 0.57 | 0.48 |
| *dsrN* | -0.52 | -0.25 | -0.16 | 0.38 | -0.57 | -0.74* | 0.37 | 0.04 | 0.04 | 0.04 | -0.08 |
| *asrA* | -0.14 | -0.25 | -0.44 | 0.29 | -0.57 | -0.74* | 0.45 | 0.04 | 0.03 | -0.17 | 0.19 |
| *cysI* | -0.10 | 0.17 | 0.58 | -0.17 | 0.31 | 0.42 | -0.39 | -0.02 | -0.03 | -0.05 | -0.60 |
| *cysNC* | 0.62 | 0.30 | 0.45 | -0.29 | 0.26 | 0.62 | -0.57 | -0.20 | -0.09 | 0.10 | 0.30 |
| *cysD* | 0.85** | 0.61 | 0.37 | -0.62 | 0.21 | 0.56 | -0.85** | -0.64 | -0.57 | -0.30 | 0.03 |
| *aprB* | 0.10 | -0.46 | -0.10 | 0.26 | -0.11 | -0.17 | 0.25 | -0.03 | 0.04 | -0.12 | 0.48 |
| *dsrO* | -0.13 | -0.87** | -0.28 | 0.33 | -0.19 | -0.37 | 0.61 | 0.49 | 0.50 | 0.45 | 0.49 |
| *dsrC* | -0.47 | -0.62 | -0.59 | 0.20 | 0.42 | -0.02 | .730* | 0.53 | 0.39 | 0.10 | -0.06 |
| *aprM* | 0.30 | -0.02 | 0.58 | 0.22 | -0.10 | 0.31 | -0.27 | 0.14 | 0.28 | 0.33 | 0.48 |
| *dsrJ* | -0.57 | -0.30 | 0.09 | 0.67 | -0.51 | -0.48 | 0.43 | 0.46 | 0.52 | 0.59 | 0.34 |
| *MET22* | 0.26 | -0.48 | 0.20 | 0.34 | 0.00 | 0.24 | 0.12 | 0.32 | 0.43 | 0.43 | 0.77* |
| *HINT4* | -0.25 | -0.53 | -0.39 | 0.24 | 0.18 | -0.07 | 0.59 | 0.58 | 0.55 | 0.43 | 0.35 |
| *SAL* | -0.03 | -0.37 | -0.48 | 0.31 | -0.27 | -0.40 | 0.49 | 0.11 | 0.09 | -0.19 | 0.32 |
| *qmoA* | -0.44 | 0.13 | -0.52 | -0.03 | 0.06 | -0.34 | 0.39 | 0.17 | 0.08 | -0.08 | -0.27 |
| *dsrD* | 0.15 | -0.25 | -.690* | 0.20 | -0.21 | -0.34 | 0.45 | -0.06 | -0.08 | -0.27 | 0.51 |

* p < 0.05, ** p < 0.01.
