## Supplementary Table S4 for "Impacts of *Desulfobacterales* and *Chromatiales* on sulfate reduction in the subtropical mangrove ecosystem as revealed by SMDB analysis"

### Supplementary Table S4. Impact of sediment properties on pathway.

| Pathways | PH | Salinity | ORP | AS | NO_3_^-^ | NH_4_^+^ | Fe | TOC | TN | TP | Sulfide |
| --- | --- | --- | --- | --- | --- | --- | --- | --- | --- | --- | --- |
| Assimilatory sulfate reduction | 0.695* | 0.26 | 0.30 | -0.54 | 0.15 | 0.33 | -0.58 | -0.37 | -0.29 | -0.15 | 0.07 |
| Dissimilatory sulfate reduction | -0.27 | -0.53 | -0.49 | 0.26 | -0.27 | -0.64 | 0.61 | 0.19 | 0.19 | -0.08 | 0.23 |
| Organic degradation/synthesis | 0.48 | 0.52 | -0.18 | -0.88** | 0.41 | 0.20 | -0.50 | -0.73* | -0.79* | -0.81** | -0.57 |
| Sulfide oxidation | 0.59 | -0.03 | 0.00 | -0.39 | 0.30 | 0.31 | -0.22 | -0.18 | -0.13 | -0.18 | 0.25 |
| Sulfite oxidation | 0.775* | 0.19 | 0.05 | -0.61 | 0.22 | 0.25 | -0.48 | -0.59 | -0.53 | -0.54 | 0.09 |
| Sulfur oxidation | -0.17 | -0.23 | -0.64 | -0.03 | -0.14 | -0.58 | 0.45 | -0.14 | -0.20 | -0.50 | -0.12 |
| Sulfur reduction | -0.36 | -0.52 | -0.66 | 0.22 | -0.20 | -0.65 | 0.704* | 0.22 | 0.14 | -0.17 | 0.02 |
| Tetrathionate oxidation | 0.15 | -0.15 | -0.03 | -0.11 | 0.41 | 0.37 | 0.07 | 0.32 | 0.31 | 0.23 | 0.15 |
| Tetrathionate reduction | -0.26 | -0.36 | -0.57 | 0.17 | -0.20 | -0.60 | 0.58 | 0.13 | 0.10 | -0.19 | 0.12 |
| Thiosulfate disproportionation | 0.51 | 0.10 | -0.01 | -0.22 | 0.08 | 0.12 | -0.20 | -0.29 | -0.22 | -0.38 | 0.28 |
| Thiosulfate oxidation | 0.679* | 0.09 | 0.05 | -0.42 | 0.38 | 0.50 | -0.33 | -0.17 | -0.13 | -0.13 | 0.26 |

p < 0.05, ** p < 0.01.
