## Supplementary Table S5 for "Impacts of *Desulfobacterales* and *Chromatiales* on sulfate reduction in the subtropical mangrove ecosystem as revealed by SMDB analysis"

### Supplementary Table S5. Sediment properties in mangrove samples and nonmangrove samples.

| Properties | MS (mean) | MS (std) | NMS (mean) | NMS (std) | p-values |
| --- | --- | --- | --- | --- | --- |
| pH | 6.76 | 0.17 | 7.03 | 0.02 | 0.049 |
| Salinity | 29.17 | 0.37 | 29.67 | 0.47 | 0.170 |
| ORP | -207.83 | 81.85 | -141.67 | 23.23 | 0.265 |
| AS | 2301.95 | 376.26 | 1383.66 | 130.34 | 0.008 |
| NO_3_^-^ | 1.18 | 0.82 | 1.60 | 1.58 | 0.664 |
| NH_4_^+^ | 15.09 | 3.39 | 20.00 | 9.01 | 0.334 |
| Fe | 96.11 | 12.52 | 54.20 | 4.25 | 0.002 |
| TOC | 11.62 | 3.73 | 3.93 | 1.07 | 0.018 |
| TN | 0.77 | 0.26 | 0.26 | 0.06 | 0.021 |
| TP | 0.15 | 0.06 | 0.10 | 0.02 | 0.272 |
| Sulfide | 0.09 | 0.05 | 0.05 | 0.01 | 0.230 |

Std: standard deviation; p-values: Student's t test of p-values.
