## Supplementary Table S6 for "Impacts of *Desulfobacterales* and *Chromatiales* on sulfate reduction in the subtropical mangrove ecosystem as revealed by SMDB analysis"

### Supplementary Table S6. Sediment properties in rhizosphere samples and nonrhizosphere samples.

| Properties | RS (mean) | RS (std) | NRS (mean) | NRS (std) | p-values |
| --- | --- | --- | --- | --- | --- |
| pH | 6.69 | 0.21 | 6.82 | 0.08 | 0.476 |
| Salinity | 29.00 | 0.00 | 29.33 | 0.47 | 0.374 |
| ORP | -142.33 | 13.91 | -273.33 | 68.01 | 0.056 |
| AS | 2583.36 | 188.85 | 2020.55 | 298.51 | 0.087 |
| NO_3_^-^ | 0.95 | 0.08 | 1.42 | 1.11 | 0.582 |
| NH_4_^+^ | 16.37 | 3.19 | 13.81 | 3.10 | 0.461 |
| Fe | 94.74 | 8.83 | 97.48 | 15.22 | 0.836 |
| TOC | 14.78 | 2.08 | 8.45 | 1.85 | 0.032 |
| TN | 1.01 | 0.12 | 0.53 | 0.07 | 0.008 |
| TP | 0.21 | 0.04 | 0.10 | 0.01 | 0.024 |
| Sulfide | 0.09 | 0.06 | 0.09 | 0.04 | 0.901 |

Std: standard deviation; p-values: Student's t test of p-values.
